# Nitroxoline-O-protected derivatives inhibit MetAP2 and activate ATF4 through mTORC1 to inhibit cancer cell growth

**DOI:** 10.1101/2025.10.20.683482

**Authors:** Michael J. Williams, Conor T. Ronayne, Tanner J. Schumacher, Kayla M. Johnson, Matthew D. Wittmer, Joseph L. Johnson, Grant W. Anderson, Venkatram R. Mereddy

## Abstract

Reprogrammed cancer cell proliferation requires high levels of protein synthesis and concomitant folding and processing. N-terminal methionine amino peptidases (MetAP) are a class of enzymes that cleave the initiator methionine amino acids to allow for peptide maturation and co-translational processing. Specifically, based on its role in protein synthesis, MetAP2 has been found to be upregulated in cancer cells and has been explored as a potential anticancer target. Cellular perturbations that impinge on protein synthesis activate cellular stress pathways, including the integrated stress response and mTORC1. Nitroxoline, a MetAP2 inhibitor has been explored as an anticancer agent but is hampered by poor pharmacokinetic properties. Here, we synthesize a few O-substituted silyl and nonsilyl nitroxoline analogs to diversify the nitroxoline template to reduce metabolic vulnerability. *In vitro* MetAP2 and cancer cell proliferation inhibition assays demonstrate that synthesized analogs retain potency when compared to the parent nitroxoline. Mechanistically, we show that the lead candidate compound **3** and nitroxoline activate ATF4 mediated stress responses through non-canonical mTORC1. These results further implicate MetAP2 protein processing in mTORC1 nutrient sensing pathways and provide novel synthetic analogs of nitroxoline for potential cancer treatment.

## INTRODUCTION

Uncontrolled cancer cell proliferation is accompanied by an increase in the demand for protein synthesis. N-terminal methionine excision (NME) is a critical co-translational process that is catalyzed by a variety of methionine amino peptidase (MetAP) enzymes that cleave the N-terminal methionine of nascent peptides as they emerge from the exit tunnel of the large ribosomal subunit^1–4^. This cleavage is necessary for the proper folding and maturation of proteins, and constitutes about two-thirds of the entire proteome^1–4^. Specifically, MetAP2 has been implicated in a variety of tumor-promoting activities including tumor growth, metastasis, and angiogenesis^5–8^. Hence, the processing of newly synthesized proteins via MetAP2 constitutes a tractable target for cancer therapy.

Inhibition of MetAP2 is predicted to result in protein folding defects. The integrated stress response (ISR) is a coordinated stress-sensing pathway that employs four kinases that converge and phosphorylate the translation initiation factor eIF2α^9^. Depending on the type of stress; mitochondrial dysfunction or heme deprivation (kinase HRI), viral infection (kinase PKR), ER-stress or protein misfolding (kinase PERK), or amino-acid deprivation (kinase GCN2), specific kinases are activated to initiate the ISR^9–11^. Phosphorylation of eIF2α inhibits global translation, and activates specific translation of specific mRNAs, including transcription factor ATF4, to activate the transcription of a variety of stress response genes^9–11^. Additionally, mTORC1 senses anabolic and catabolic states of the cell, and is known to modulate the expression of ATF4^12–14^. Thus, mTORC1 is also a predicted mediator of cellular stress accompanied by defects in protein processing through MetAP2 inhibition.

Nitroxoline (5-nitroquinolin-8-ol) is an antibiotic classically used to treat urinary tract infections. Interestingly, nitroxoline also exhibits potent MetAp2 inhibition properties^7,8^. The anticancer efficacy of nitroxoline was investigated in *in vivo* human and murine mouse tumor models. These studies indicated that nitroxoline reduced tumor growth in both xenograft and syngraft models^5–8^. However, anticancer applications of nitroxoline therapy pose significant constraints based on unfavorable pharmacokinetic properties^7^.The metabolic liability of nitroxoline is mainly the free phenol which goes through rapid sulfation and is quickly eliminated^7^. As a result, substantial doses are needed to suppress tumor growth limiting the possibility for clinical translation for anticancer applications. Protecting the phenol group with labile and non-labile small protecting groups should stabilize the molecule from glucuronidation and lead to a longer duration of the drug in the blood to get to the target site.

In the current work, we synthesized and evaluated novel synthetic analogs of nitroxoline as anticancer agents. Here, by appending the unprotected phenol of nitroxoline with a variety of silyl groups we hypothesize that these drug candidates will have enhanced pharmacokinetic properties when compared to the parent nitroxoline, as silyl groups are lipophilic and are cleaved only under acidic conditions^15^. Lead candidates retained MetAP2 inhibition properties, and were found to inhibit MetAP2 with a conserved non-competitive mechanism, *in vitro*. Further, cancer cell proliferation studies revealed that silyl derivatives retained similar activity to the parent nitroxoline. Finally, we illustrate that MetAP2 inhibition with lead candidate compounds activates the ISR evidenced by marked increases in ATF4 expression, independent of eIF2α phosphorylation. Treatment with mTORC1 inhibitor, rapamycin, reversed MetAP2 inhibitor induced ATF4 expression, implicating this nutrient sensing pathway in the cellular stress response. These results further diversify the synthetic template of nitroxoline and provide insights into the cellular stress mechanisms that impinge on MetAP2.

## RESULTS

### Design and synthesis of novel O-substituted silyl and nonsilyl nitroxoline derivatives with potential for improved PK properties

The pharmacological activity of nitroxoline (**1**) has been largely attributed to its MetAP2 inhibition characteristics. However, the anticancer utility of this molecule has been hampered by limited metabolic stability^7^. Specifically, it is well characterized that the phenol group of nitroxoline is particularly vulnerable to phase-two metabolism (**Figure 1A**). In this regard, we synthesized a variety of silyl and non-silyl substituted lipophilic nitroxoline derivatives **2**-**9** to protect this metabolic vulnerability of the unprotected phenol group of nitroxoline (**Figure 1B**). We evaluated the MetAP2 inhibition and cellular proliferation inhibition properties of the synthesized compounds to identify a lead candidate MetAP2 inhibitor.

**Figure 1.**
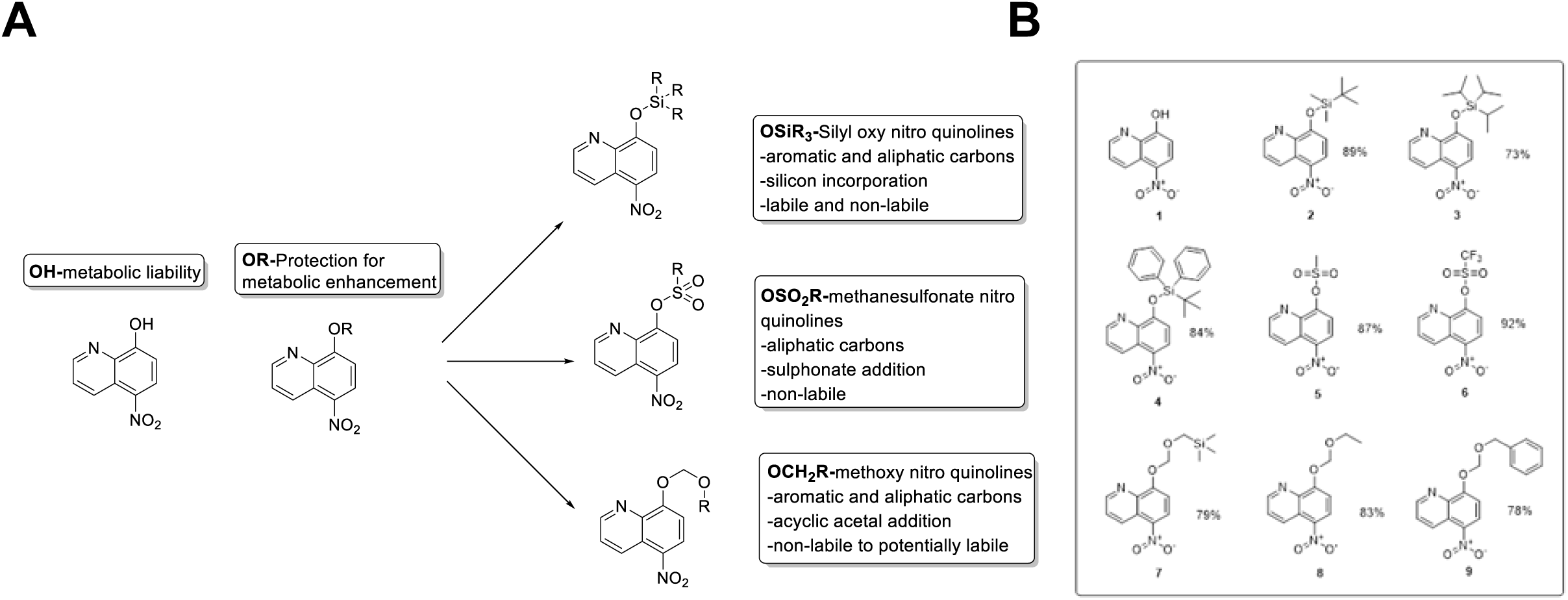
(**A**) Schematic highlighting the metabolic vulnerability of nitroxoline -OH and the synthetic modifications imparted on this template. Note the variety of silyl and non silyl groups and their characteristics. (**B**) Synthesized nitroxoline analogs and their respective yields.

### Synthetic silyl and non-silyl nitroxoline analogs **2-9** retained MetAP2 inhibition properties of parent nitroxoline **1**

To investigate the ability of compounds **2-9** to inhibit MetAP2 activity, we employed an *in vitro* fluorometric assay using recombinant MetAP2 and substrate methionine-aminocoumarin (H-Met-AMC), which fluoresces upon enzymatic hydrolysis (**Figure 2A**). Fluorescence generated by AMC release in the presence of MetAP2 reports on enzymatic activity and decreases in fluorescence in the presence of compounds provides a means to evaluate the inhibitory capacity of synthesized compounds. In this regard, we tested parent nitroxoline, and synthesized derivatives **2-9**, in a dose response fashion (ex. **Figure 1B**), to determine their IC_50_ value of MetAP2 activity. Here, we found that compound **2** (**Figure 2B, E**), along with other silyl derivatives **3** and **4** (**Figure 2C-D, E**) retained MetAP2 inhibition characteristics. Interestingly, trifluouromethyl-silyl oxide compound **6**, and non-silyl protecting groups on compounds **7**-**9** lost activity, illustrating the necessity of the chemical nature of the silyl groups in compounds **2**-**4** (**Figure 2E**). Further, Michaelis–Menten kinetics and Lineweaver Burk plots revealed that test compounds inhibited MetAP2 in a similar non-competitive fashion when compared to parent nitroxoline **1** (**Figure 2F-G**). These results illustrated that the synthetic modifications on the candidate compounds **2-4** did not lose MetAP2 inhibition characteristics for further investigation.

**Figure 2.**
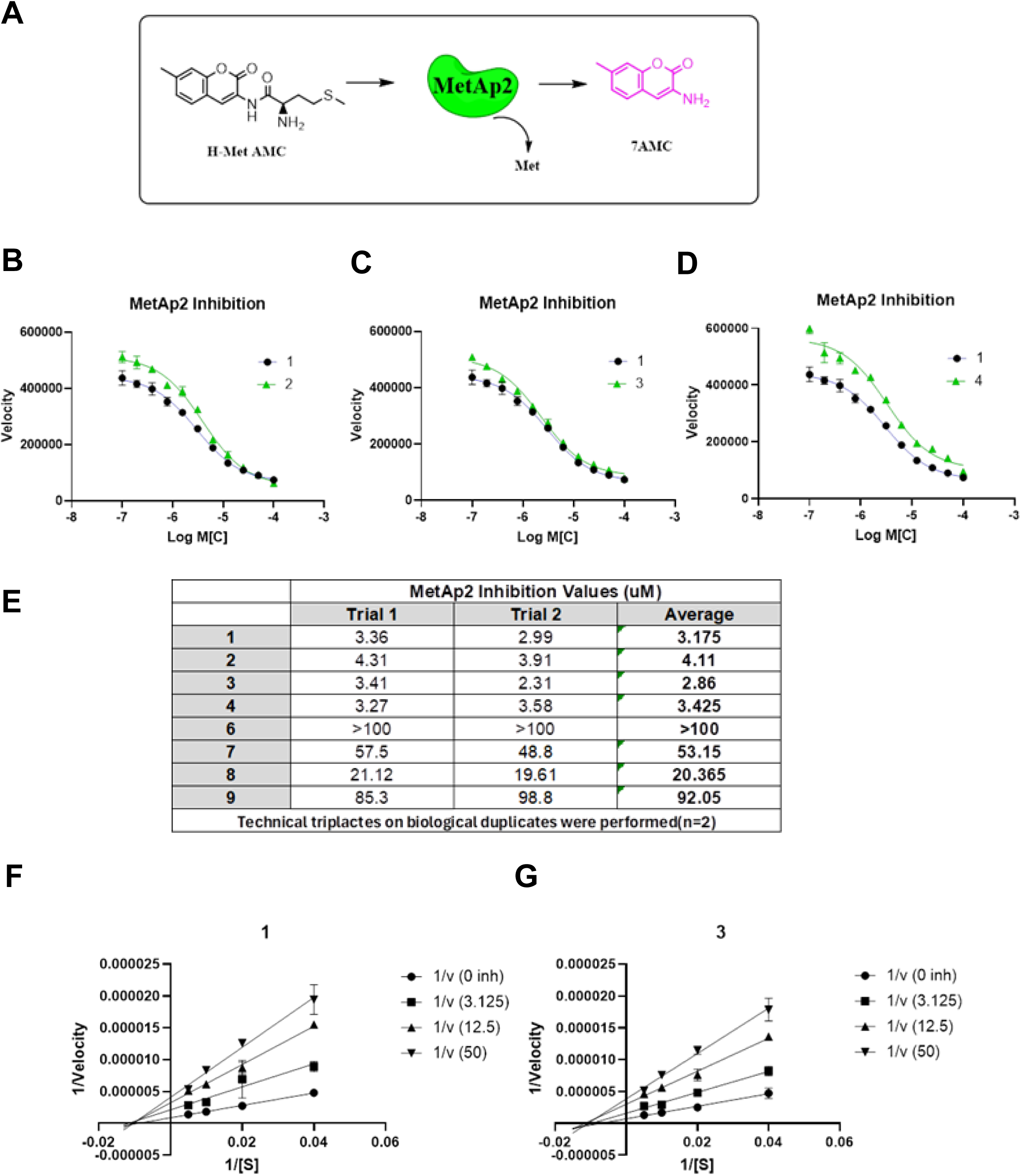
(A) Biochemical scheme of *in vitro* recombinant MetAP2 inhibition assay. Non-fluorescent MetAP2 substrate H-Met-AMC is hydrolyzed by MetAP2 resulting in fluorescent 7AMC molecule. Note methionine cleavage in this reaction. (B-D) MetAP2 inhibition characteristics and dose response curves compared to nitroxoline 1 and (B) compound 2, (C) compound 3, and (D) compound 4. (E) Respective compounds average IC_50_ (µM) values two independent experiments (n=2). (F) Lineweaver Burk plot for nitroxoline **1** with the method described above, eight inhibitor concentrations with five substrate variations. (N=2, n=2). (G) Lineweaver Burk plot for compound **3** with the method described above, eight inhibitor concentrations with five substrate variations. (N=2, n=2).

Previous reports indicate that nitroxoline can also inhibit Cathepsin B, a lysosomal cysteine protease. In a similar *in vitro* enzymatic assay, recombinant Cat B and fluorescent substrate Z-Leu-Arg-AMC were used to screen the inhibitory capacity of nitroxoline and synthetic derivatives, where inhibition of Cat B was only observed at high µM concentrations, indicating that MetAP2 remains a primary target (appendix, data not shown).

### Synthetic nitroxoline analogs **2-9** retained anti-cancer cell proliferation properties of parent nitroxoline **1**

MetAP2 inhibitors have exhibited promising anticancer efficacy in advanced solid tumors^5–8^. In this regard, we evaluated the *in vitro* cell proliferation inhibition of our candidates against a panel of human (MDA-MB-231) and mouse (67nr and 4T1) breast cancer and human pancreatic cancer (MIAPaCa-2 and Bx-PC3) cell lines. MetAP2 expression levels were confirmed in these cell lines, where we noted relative steady state MetAP2 protein levels varied across these selected cell lines (**Figure 3A**). To evaluate cancer cell proliferation inhibition properties, we treated respective cancer cell lines with test compounds **1-9** in a dose dependent fashion for 72 hours, after which we evaluated cell viability using the widely employed MTT assay. Using untreated cells as controls, we calculated IC_50_ values of each compound as the concentration of test compound to inhibit cancer cell growth to 50% of controls (**Figure 3B**). Here, we observed that test compounds **2-9** largely retained the potency of cancer cell inhibition properties when compared the parent nitroxoline **1**, with a few exceptions. Interestingly, the potency of test compounds did not directly correlate with MetAP2 expression status in these cell lines, suggesting that cell survival in the context of MetAP2 may be dependent on intermediary and regulatory signaling mechanisms that may differ between cell lines. Nonetheless, these results indicated that the silylation of nitroxoline phenolic OH does not alter the cellular treatment responses (**Figure 3B**), which is consistent with retained MetAP2 inhibition efficiency (**Figure 2**). Specifically, we have selected compound **3** as a lead compound to compare with parent nitroxoline **1** based on its retained MetAP2 inhibition and slightly enhanced cellular potency.

**Figure 3.**
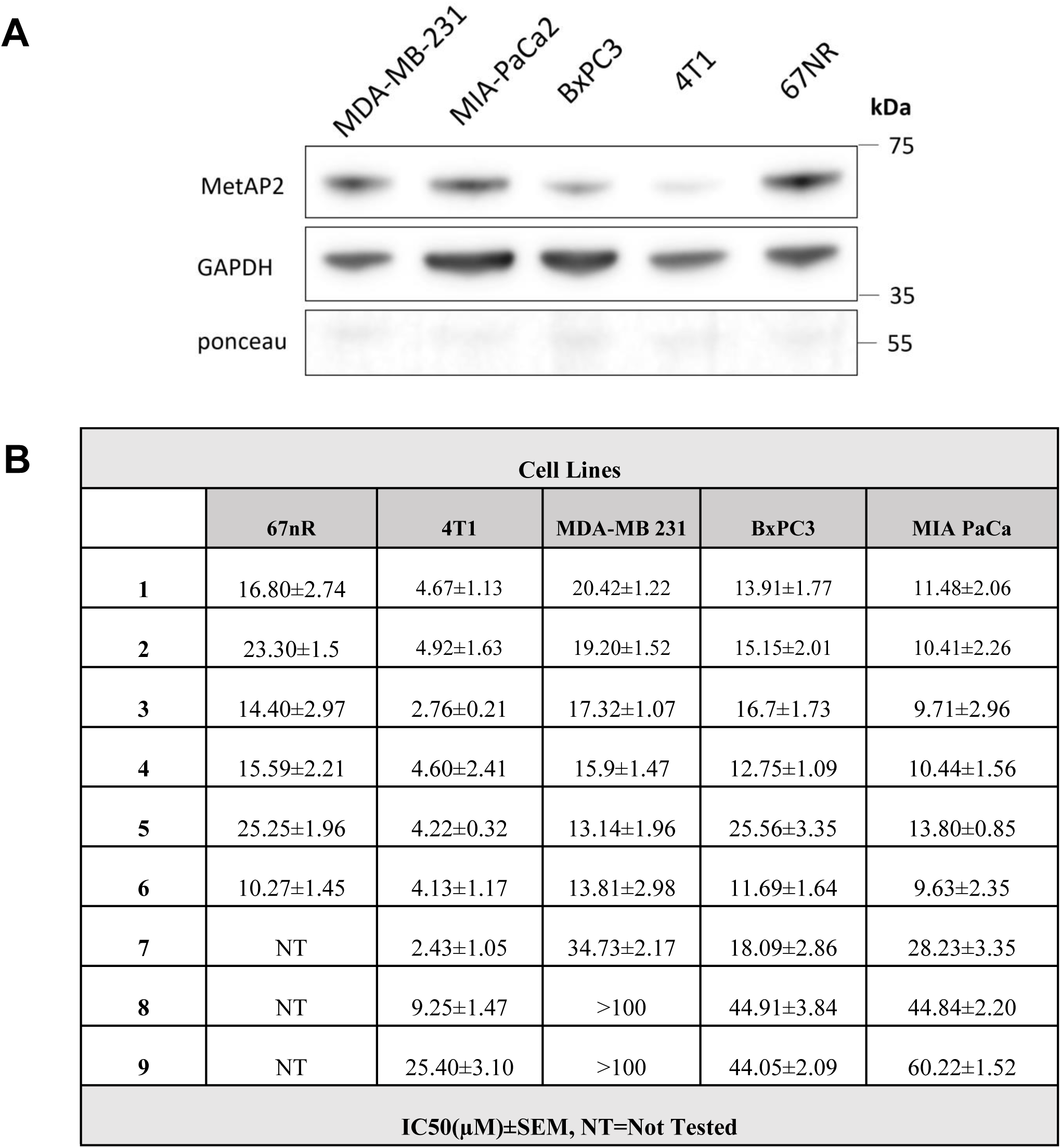
(**A**) MetAP2 expression levels on tested cell lines MDA-MB-231, MIAPaCa-2, BxPC-3, 4T1, and 67NR. (**B**) Cellular IC_50_ values of candidate compounds against the indicated cell lines. Values represent the average ± SEM of at least three independent experiments (N=3).

### MetAP2 inhibition with synthetic silyl nitroxoline analog **3** activates ATF4 through mTORC1

MetAP2 inhibition results in defects in protein processing and is hypothesized to activate cellular stress pathways involved in protein homeostasis. Specifically, the canonical integrated stress response (ISR) responds to defects in protein homeostasis largely through the activation of ISR kinases that converge on the phosphorylation of translation initiation factor eIF2α to promote specific translation of the transcription factor ATF4^9^. In this regard, we investigated the ability of candidate compound **3** to promote the expression of ATF4 in murine (**Figure 4A**) and human (**Figure 4B**) breast and pancreatic cancer cells. Interestingly, in the murine breast cancer cell lines, nitroxoline **1** and candidate **3** induce ATF4 expression in 67nr, and not in 4T1 (**Figure 4A**). Notably, 67nr exhibits marked increased MetAP2 expression when compared to 4T1, and the ability of candidate MetAP2 inhibitors to induce ATF4 expression correlates with basal MetAP2 expression levels. These observations extended to human cancer cell lines MDA-MB-231, MIAPaCa-2, and Bx-PC3 cell lines (**Figure 4B**). Importantly, the increases in ATF4 expression levels in compound treated cultures appeared to be independent of the canonical phosphorylation of eIF2α (**Figure 4A&B**). These results suggested that ATF4 was being activated through a separate mechanism. It is well documented that ATF4 can also be activated via mTORC1, a master regulator of anabolic and catabolic states of the cell mainly through a balance of protein synthesis and autophagy^12^. Hence, we hypothesized that protein dysfunction as a result of MetAP2 could be sensed by mTORC1 to activate ATF4 mediated stress responses. To test this, we assessed the ability of mTORC1 inhibitor, rapamycin, to reverse the nitroxoline **1** and compound **3** induced ATF4 expression. Here, we observed that MDA-MB-231(**Figure 4C**) and MIAPaCa-2 (**Figure 4D**) cells treated with test compounds resulted again in increased expression of ATF4 which was reversed by rapamycin (**Figure 4C&D**). These results strongly support the notion that MetAP2 inhibition is communicating stress signals through mTORC1 to activate ATF4 expression.

**Figure 4.**
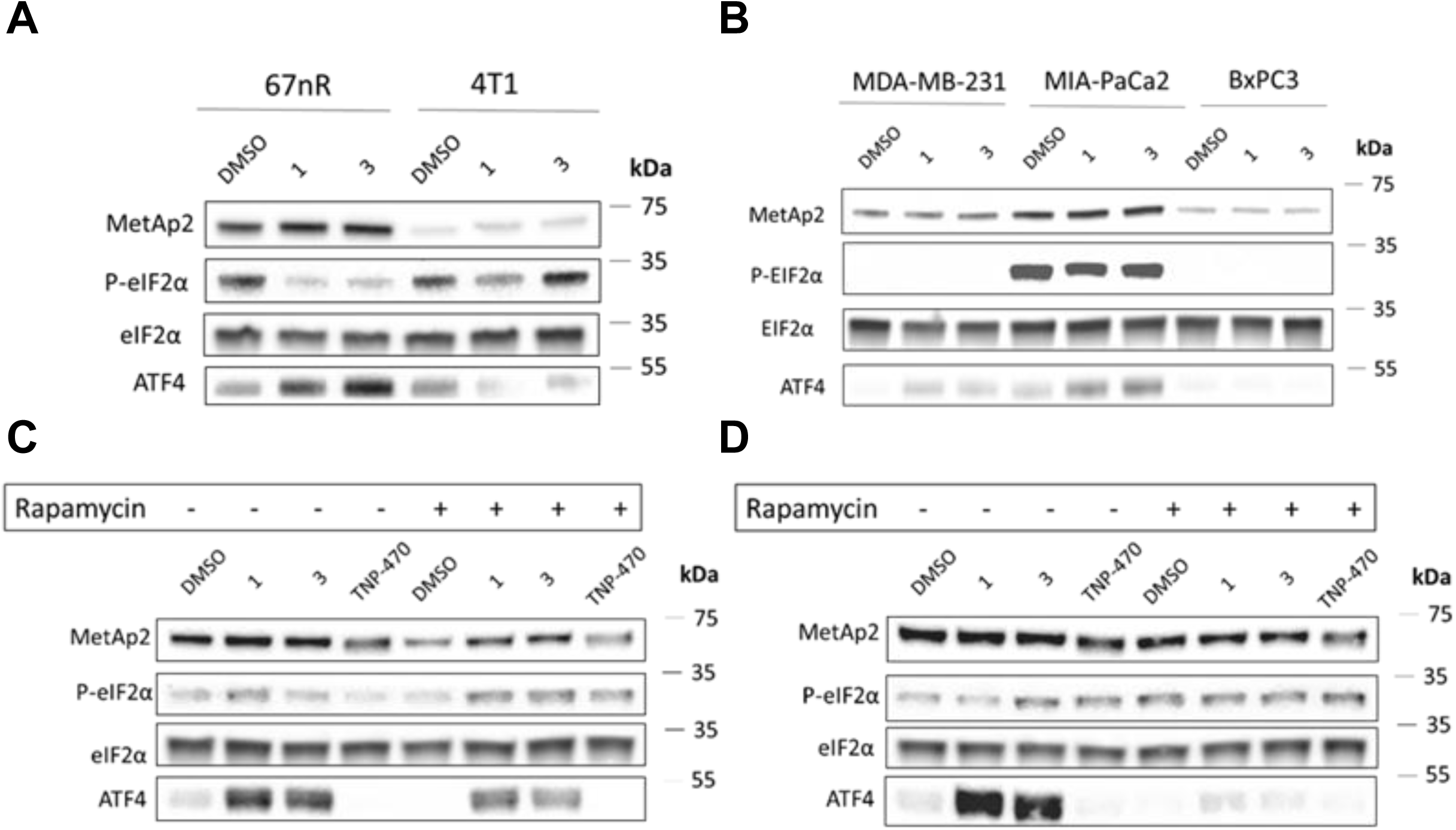
(**A**) Murine cells 67nR and 4T1 treated with DMSO, Nit, and Nit-OTIPS (N=3, n=2) (**B**) Human derived cell lines MDA-MB-231, Mia PaCa2, BxPC3 treated with DMSO, Nit, and Nit-OTIPS (N=3, n=2), (**C**) MDA-231 cells treated with DMSO, Nit, Nit-OTIPS, and TNP-470. The same dosing was repeated in combination with 50nM rapamycin denoted as the “+” groups. (N=3, n=2). (**D**) Mia PaCa2 assays, cells were treated with DMSO, Nit, Nit-OTIPS, and TNP-470. The same dosing was repeated in combination with 50nM rapamycin denoted as the “+” groups. (N=3, n=2).

## DISCUSSION

Many pharmacologically active drugs fail to make it to the clinic due to unfavorable pharmacokinetic and pharmacodynamic properties. Although nitroxoline significantly reduces tumor growth in multiple tumor models, it has metabolically unfavorable properties mainly due to a phenol group. Since the pH of cancer cells is mildly acidic (pH 6.3-7.0) relative to physiological pH (7.1-7.3), it was hypothesized that introducing labile and non-labile functional groups to the phenolic OH would prevent the sulfation step, enhancing metabolic stability. In addition, a lower pH inside the cancer cells could cleave the silyl-protecting group (**2-4**) to release the parent nitroxoline. This should provide anticancer efficacy at a clinically relevant dose. Even if the silyl protecting groups are not cleaved inside the cancer cells, they should still be valuable derivatives if they retain similar cell proliferation properties and the same mechanism of action as the parent nitroxoline. Apart from silyl derivatives (**2-4**), non-cleavable triflate and mesylate derivatives (**5-6**), and moderately cleavable candidates (**7-9**) were synthesized.

Using compounds **1-9**, cell proliferation inhibition concentrations (µM) on cell lines 67nR, 4T1, MDA-MB-231, BxPC3, and MIA PaCa were determined. Compounds **2-4** showed a ∼15% increased efficacy across all cell lines except BxPC3, which was equal to 1. Cell proliferation efficacy was reduced nearly twofold for compound **5** on 67nR, MDA-MB 231, BxPC3, and MIA PaCA but was enhanced ∼10% in 4T1. Compound **6** displayed enhanced cell proliferation potency compared to **1** on all cell lines. Cell proliferation was reduced nearly two fold for the same concentration of nitroxoline for compound **5** on 67nR, MDA-MB 231, BxPC3, and MIA PaCA but was enhanced ∼10% in 4T1. Compounds **7-9** displayed reduced cell proliferation values on all cell lines except for **7**, showing a two-fold increase in 4T1. Silyl modifications to nitroxoline (**2-4**) maintained or slightly enhanced efficacy. When a non-labile mesylate (**5**) protecting groups were added, there was no benefit in cell proliferation inhibition. Compounds **7-9** showed reduced efficacy, which could be justified by the addition of a second oxygen in the protecting chain coming on the phenol of nitroxoline, a more significant chemical addition, and therefore a different MOA.

Inhibition of MetAp2 exhibited equipotency for compounds **3** and **4** to that of the control, nitroxoline, **1** (∼3.4µM), and compound **2** a 21% reduction in inhibition efficacy (4.3µM). Compound **7** showed margin inhibition at 21.1 µM, while the chemical modifications in **6**, **8**, and **9** drastically reduced the IC_50_ resulting in near if not over 100 µM, making them non potent. Based on this data including lipophilicity, cell proliferation inhibition efficacy, and MetAP2 inhibition, compound **3** was chosen as a lead candidate. To characterize the type of inhibition nitroxoline and compound **3** were displayed as Lineweaver-Burk plots for each were conducted (n=3). Nitroxoline exhibited that at different concentrations the lines merge at x=0, while the y-intercept changes, these results suggest that noncompetitive inhibition of MetAp2 is occurring with the treatment of nitroxoline. Compound **3** also exhibited that at different concentrations lines merge at x=0, while the y-intercept changes, suggesting that noncompetitive inhibition of MetAp2 is occurring with nitroxoline-OTIPS. These results indicate that TIPS protection to the phenol on nitroxoline results in the same type of inhibition as the parent drug, nitroxoline.

Investigations on the effect of treatment with nitroxoline and compound **3** on the well-defined key players involved in the ISR blotting were performed on murine and human cell lines. The murine cancer model 67nR, with the treatment of nitroxoline and compound **3**, showed minimal p-eIF2α present. To our surprise, with drug treatment, ATF4 levels were enhanced, while p-eIF2α decreased, hinting at a non-canonical response to the well-defined pathway. The murine cancer model 4T1, with the treatment of nitroxoline and compound **3**, showed no expression changes compared to that of the control with any protein probed. With compound treatment, ATF4 levels were diminished, while p-eIF2α was present, but band intensity was homogeneous for all treatment groups. The investigations into the effects of nitroxoline and compound **3** on human cancer began with the breast cancer cell line MDA-MB-231. The results highlighted a similar phenomenon occurring in 67nR, the non-canonical activations of ATF-4 with drug treatment. Furthermore, in two human pancreatic cancer models, MIA PaCa and BxPC3, the effects of treatment with nitroxoline and compound **3** were determined. MIA PaCA drug treatment led to the highest activation across all cancer cell lines studied. In MIA PaCa, the presence of p-eIF2α was apparent but equal across treated and non-treated groups, showing cell-specific characteristics and non-canonical activation of ATF4.

## CONCLUSIONS

In conclusion we have designed, synthesized, and evaluated novel O-substituted silyl and non-silyl nitroxoline analogs as MetAP2 inhibitors. Here, we illustrated that silyl groups did not decrease the MetAP2 inhibition or cancer cell proliferation inhibition properties. Interestingly we identified that these compounds, activate the integrated stress response as evidenced by ATF4 expression, and did so via non-canonical mTORC1. These studies provided a means to potentially overcome the pharmacokinetic shortcomings of nitroxoline, and illustrated downstream stress signaling involves mTORC1 and ATF4.

## METHODS

### Materials and equipment

Nitroxoline, imidazole, tert-butyldimethyl silyl chloride, triisopropylsilyl chloride, tert-butyl (chloro)diphenyl silane, methane sulfonyl chloride, trifluoromethanesulfonic anhydride, 2- (trimethylsilyl) ethoxymethyl chloride, 2-methoxyethoxymethyl chloride, and benzyl chloromethyl ether were purchased from Ambeed (Arlington Heights, IL). 1 H and 13C NMR spectra were determined using a Bruker Ascend™ 400 spectrometer. HRMS were determined using a Bruker micrOTOF-Q III ESI mass spectrometer. RFU output and absorbance values were recorded using a BioTek SYNERGY, neo2, multi-mode reader (version 3.09) with Gen5 Microplate Reader and Imager Software (BioTek Instruments Inc., Winooski, VT). Immunoblots were imaged using a Licor imager with Licor Image Software 6.0 (LI-COR Biotechnology, Lincoln, NE).

### Statistical analysis

Statistical analysis was conducted using GraphPad Prism 10 software (GraphPad Software, Boston, MA, USA). In vitro assays were performed independently in triplicate with DMSO as a control. Data was analyzed using Student’s t-test and ANOVA and log transformation was used when appropriate. P values <0.05 were considered statistically significant.

### Synthesis of silyl(oxy)-5-nitroquinolines: Compounds **2-4**

8-Hydroxy-5-nitroquinoline (Nitroxoline) 1 (1.0 mmol) was dissolved in DCM and cooled to 0°C. Imidazole (3.0 mmol) was added and stirred for five minutes. Each representative silyl chloride; tert-butyldimethyl silyl chloride, triisopropylsilyl chloride, tert-butyl chlorodiphenyl silane (1.2mmol) was individually added dropwise over five minutes, the reaction vessel was capped and under continuous stirring allowed to come to room temperature and further reacted for 1-3 hours. TLC was used to monitor reaction progress and verify reaction completion. (30% EtOAc/Hex) DCM then evaporated, and the crude oil was worked up with H_2_O/EtOAc. The ethyl acetate layer was then dried using MgSO_4_, and after filtering, the product was concentrated using a rotary evaporator. The reaction product was crude, so column chromatography used 10% EtOAc/hexanes and was eluted with DCM. Recrystallization using hexane was performed, providing adequate yields (73-89%) generating compounds **2-4**.

### Synthesis of triflate and mesylate derivatives of nitroxoline **5** and **6**

Nitroxoline, **1** (1.0 mmol) was dissolved in DCM and cooled to 0°C. Pyridine (3.0 mmol) was added and stirred for five minutes Methane sulfonyl chloride or trifluoromethanesulfonic anhydride (1.2 mmol) was added dropwise over five minutes, the reaction vessel was capped and under continuous stirring allowed to come to room temperature and further reacted for 1 hour. TLC was used to monitor reaction progress and verify reaction completion. (30% EtOAc/Hex) DCM was then evaporated, and the crude oil was worked up with H_2_O/EtOAc. The ethyl acetate layer was then dried using MgSO_4_, and after filtering the product was concentrated using a rotary evaporator. The crude product was then further purified using column chromatography (10% EtOAc/hexanes, eluted with DCM) and hexane recrystallization gave adequate yields (87, and 92%) generating compounds **5** and **6**.

### Synthesis of 8-(ethoxymethoxy)-5-nitroquinoline derivatives **7-9**

Nitroxoline 1 (1.0 mmol) was dissolved in DCM and cooled to 0°C. Imidazole (3.0 mmol) was added and stirred for five minutes. Ethoxy chlorides (1.2 mmol) were added dropwise over 5 minutes, the reaction vessel was capped and under continuous stirring allowed to come to room temperature and further reacted for 2-6 hours. TLC was used to monitor reaction progress and verify reaction completion. (30% EtOAc/Hex) DCM was then evaporated, and the crude oil was worked up with H_2_O/EtOAc. The ethyl acetate layer was then dried using MgSO_4_, and after filtering, the product was concentrated using a rotary evaporator. The crude product was then further purified using column chromatography (10% EtOAc/hexanes, eluted with DCM) and hexane recrystallization gave adequate yields (78-83%) generating compounds **7-9**.

### Spectra analysis

See appendix for spectra from compounds **2-9**.

### MTT Cell Proliferation Assay

Cell proliferation inhibition properties were evaluated using the MTT assay. Cancer cells were seeded in 96 well plates (5x103 cells/well) and incubated for 12-24 hours. Test compounds were added to respective cell lines in a serial dilution fashion for a total of eight concentrations. DMSO was used to dissolve 100mM stock solutions at 1000X starting concentration, resulting in DMSO cell concentration of 0.1% (v/v). Cells were incubated in a test compound for 72 hours. 3-(4,5-dimethylthiazol-2-yl)-2,5- diphenyltetrazolium bromide (MTT, 0.5 mg/mL) was added and incubated with the cells for four hours at 37°C under 5% CO2. To ensure formazan precipitate the solid was dissolved in a sodium dodecyl sulfate solution (0.1 g/L 0.01 N HCL) and incubated for four hours. Absorbance readings at 570 nm were recorded. IC50 values of compounds were determined using the absorbance of untreated wells as the control value for cell survival measurements.

### Human recombinant Methionine Aminopeptidase 2 screening assay

Human recombinant Methionine Aminopeptidase 2 (MetAp2) was purchased from R&D Systems, bio-techne. The fluorogenic substrate H-Met-AMC was purchased from Chemodex. Reaction buffer contained100mM Tris-maleate, 0.1mM CoCl2, 100mM NaCl, 0.05% BSA, pH=7.5. Nitroxoline (8-hydroxy-5-nitroquinoline) was purchased from Millipore Sigma. The enzymatic reaction was performed using a Costar 96 well black opaque half-well. Fluorescence intensity was recorded using a BioTek SYNERGY, neo2, multi-mode reader (version 3.09) with Gen5 Microplate Reader and Imager Software. The assay was performed at 37°C using Kinetic mode for 1 hour reading each well every 21 seconds. (λexcitation=360 nm; λemission=460nm) The production of the fluorogenic product, 7-methyl coumarin, was determined by relative fluorescent units (RFU) over time.

### Human Recombinant Methionine Aminopeptidase 2 Inhibitor Screening Assay

The test compounds and the vehicle (DMSO) were tested in triplicate. Screening concentrations began at 100µM and were further serial diluted 8 times. The positive control nitroxoline was tested on the same plate in triplicate at the same concentration as the test compounds. The plate was incubated with compounds at 37°C for 10 minutes, adequate time for binding. A 1:1 dilution was performed using a multichannel pipette in all test wells using the substrate dissolved in the buffer to achieve a final concentration of 50µM was added activating the enzymatic reaction uniformly. RFU output was recorded using a BioTek SYNERGY, neo2, multi-mode reader (version 3.09) with Gen5 Microplate Reader and Imager Software. The assay was performed at 37°C using Kinetic mode for 1 hour reading each well every 21 seconds. (λexcitation=360 nm; λemission=460nm) The production of the fluorogenic product, 7-methyl coumarin was determined by relative fluorescent units (RFU) over time. The RFU values were plotted for each concentration and normalized to the negative control, 50µM substrate with the vehicle. The triplicate samples were plotted as IC50 values, generated for nitroxoline and each novel compound (Graphpad Prism 10.0).

### Immunoblotting: MetAp2, ATF-4, eIF2α total, and phospho-eIF2α

Cancer cell lines were cultured in RPMI media containing 5%FBS and 0.5% penicillin-streptomycin, seeded at 5.0x105 cells per 60mm dish and incubated for 24 hours at 37°C under 5% CO_2_. Compounds along with (DMSO)were added to the dishes at a concentration of 50% of the predetermined IC_50_ value to initiate a response at a non-lethal dose. The dishes were incubated at 37°C under 5% CO_2_ for an additional 24 hours. Post-treatment the media was aspirated and washed two times with cold PBS. Dishes were placed on ice and stored at -85°C. 50µl of a 1X RIPA solution containing protease and phosphatase was added to each plate. The dishes were scraped, and the mixture pipetted into a 1 ml Eppendorf. Each sample was pulsed for 4 seconds with a probe sonicator and centrifuged at 0°C. BCA standardization was performed on all samples for homogenesis loading. The protein lysates were mixed with water and 5X Laemmli with 10% B-mercaptoethanol and stored at -85°C.

## APPENDIX

### Cathepsin B screening assay

Human recombinant Cathepsin B was purchased from BPS Bioscience (San Diego, CA, USA). The fluorogenic Cathepsin V substrate (Z-Leu-Arg-AMC) used for Cathepsin B was also provided by BPS Bioscience, along with the company’s proprietary Cathepsin buffer. E-64 is an irreversible and selective cysteine protease inhibitor. i.e., Cathepsin inhibitor. (BPS Bioscience) The enzymatic reaction was performed using a Costar 96 well black opaque half-well. Fluorescence intensity was recorded using a BioTek SYNERGY, neo2, multi-mode reader (version 3.09) with Gen5 Microplate Reader and Imager Software. The assay was performed at 21°C using Kinetic mode for 1 hour reading each well every 21 seconds (λexcitation=360 nm; λemission=460nm). The production of the fluorogenic product, 7-methyl coumarin was determined by relative fluorescent units (RFU) over time.

### Human recombinant cathepsin B inhibitor screening assay

Each test well contained 0.01ng of human recombinant Cathepsin B and was diluted accordingly with the provided buffer. The test compounds and the vehicle (0.05%DMSO) were tested in triplicate. Screening concentrations began at 500µM and were further serial diluted 12 times. The positive control E-64, a known suicide inhibitor of Cathepsin B was tested on the same plate in triplicate at a final concentration of 0.01µM. A 1:1 dilution was performed using a multichannel pipette in all test wells using the substrate dissolved in the buffer to achieve a final concentration of 5µM, activating the enzymatic reaction uniformly. RFU output was recorded using a BioTek SYNERGY, neo2, multi-mode reader (version 3.09) with Gen5 Microplate Reader and Imager Software. The assay was performed at 21°C using Kinetic mode for 1 hour reading each well every 21 seconds. (λexcitation=360 nm; λemission=460nm) The production of the fluorogenic product, 7-methyl coumarin was determined by relative fluorescent units (RFU) over time. The RFU values were plotted for each concentration and normalized to the negative control, 5µM substrate with the vehicle. The triplicate samples were plotted as IC50 values, generated for nitroxoline and each novel compound (Graphpad Prism 10.0).

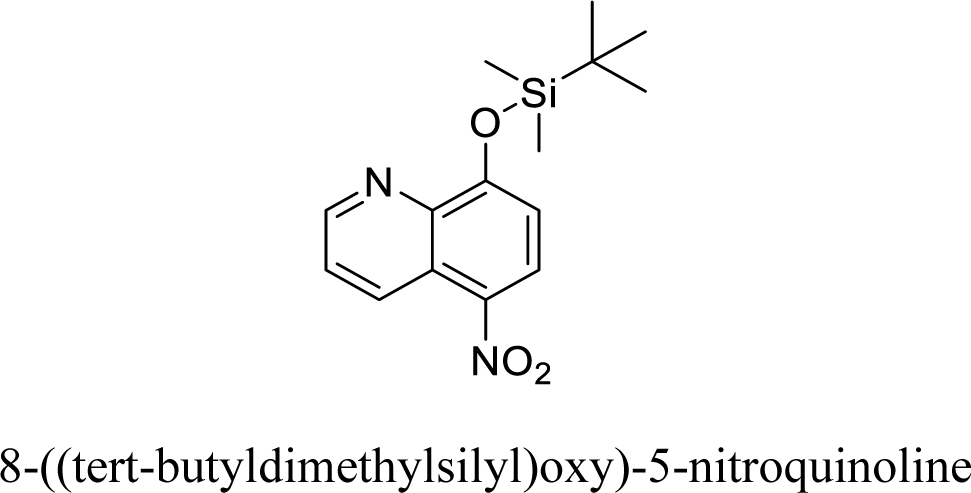

**^1^H NMR (400 MHz, Dimethyl Sulfoxide-d6):** δ 9.19 (d, J= 8.88 Hz 1H), 9.05, 8.58 (d, J= 8.8 Hz, 1H), 7.92 (dd, J=4.12 Hz, 1H), 7.24 (d, J= 8.84 Hz, 1H), 0.88 (s, 9H), 0.00 (s, 6H)

**^13^C NMR (100 MHz, Dimethyl Sulfoxide-d6):** δ 163.36, 151.90, 140.01, 137.82, 135.12, 131.76, 127.94, 125.25, 112.71, 28.50, 20.49, -0.64

**HRMS (ESI) m/z:** Calculated for C_15_H_20_N_2_O_3_Si+H^+^: 304.1216, found 304.1234

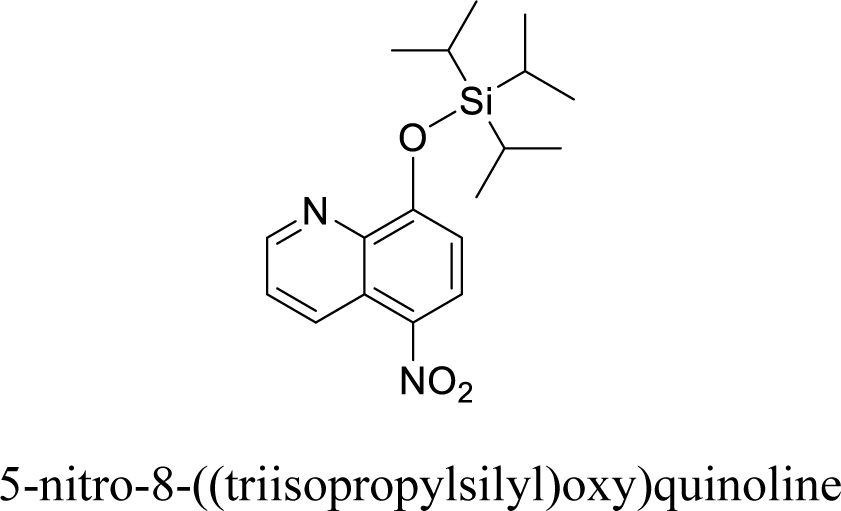

**^1^H NMR (400 MHz, Dimethyl Sulfoxide-d6):** δ 9.15 (d, J= 10.3 Hz 1H), 9.02 (d, J=4.12,1H), 8.55 (d, J= 8.84 Hz, 1H), 7.88 (dd, J=4.12 Hz, 1H), 7.20 (d, J= 8.84 Hz, 1H), 1.02-0.08 (m, 21H)

**^13^C NMR (100 MHz, Dimethyl Sulfoxide-d6):** δ 161.12, 149.61, 137.75, 135.54, 132.85, 129.50, 125.67, 123.00, 110.44, 18.27, 12.53

**HRMS (ESI) m/z:** Calculated for C_18_H_26_N_2_O_3_Si+H^+^: 347.1785, found 347.1774

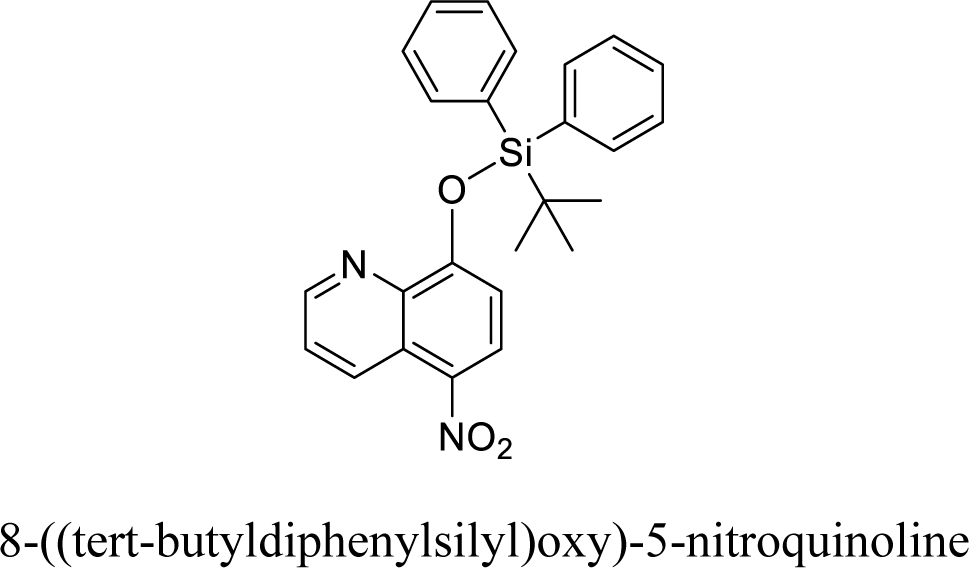

**^1^H NMR (400 MHz, Dimethyl Sulfoxide-d6):** δ 9.16 (m, 1H), 9.02 (m, 1H), 8.55 (d, J=8.84 Hz, 1H), 7.88 (m, 1H), 7.71 (q, 4H), 7.40 (m, 6H), 7.21 (d, J=8.84Hz, 1H), 0.97 (s, 9H)

**^13^C NMR (100 MHz, Dimethyl Sulfoxide-d6):** δ 161.12, 149.64, 137.77, 136.84, 135.57, 134.95, 132.87, 129.63, 129.52, 127.96, 125.69, 123.01, 110.46, 27.01, 19.14 ppm

**HRMS (ESI) m/z:** Calculated for C_25_H_24_N_2_O_3_Si+H^+^: 429.1629, found 429.1642

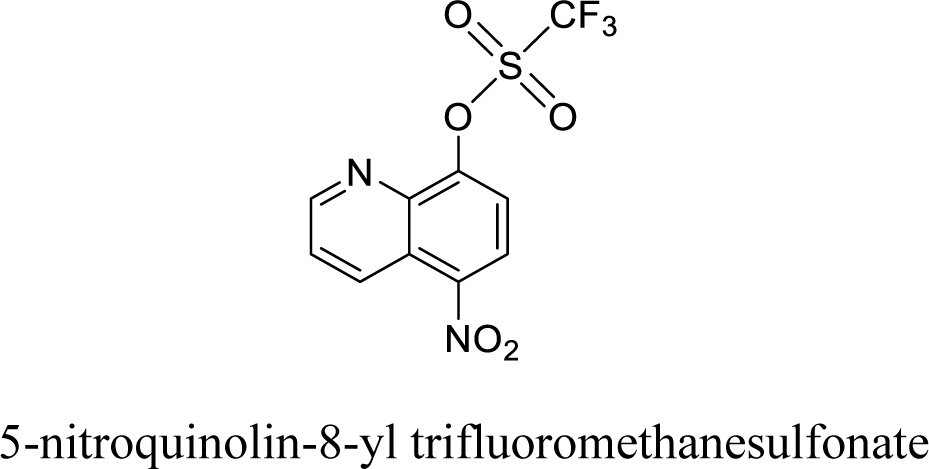

**^1^H NMR (400 MHz, Dimethyl Sulfoxide-d6):** δ 9.23 (d, J= 3.4 Hz 1H), 8.95 (dd, J=8.8,1H), 8.58 (d, J= 8.6 Hz, 1H), 8.15 (d, J=8.56 Hz, 1H), 7.98 (q, J= 8.88 Hz, 1H)

**^13^C NMR (100 MHz, Dimethyl Sulfoxide-d6):** δ 153.52, 148.89, 145.23, 140.25, 132.85, 126.14, 125.64, 123.55, 122.49, 121.11, 120.37, 117.18, 114.00

**HRMS (ESI) m/z:** Calculated for C_10_H_5_F_3_N_2_O_5_S+H^+^: 321.9944, found 321.9941

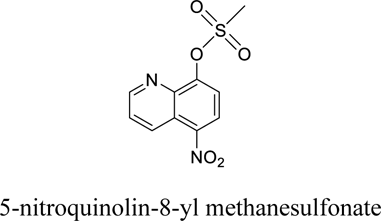

**^1^H NMR (400 MHz, Dimethyl Sulfoxide-d6):** δ 9.19 (d, J= 4.08 Hz 1H), 8.93 (d, J= 8.84 Hz, 1H), 8.56 (dd, J=8.56,1H), 8.00 (q, J=8.52 Hz, 1H), 7.97-7.92 (m, 1H), 3.66 (s, 3H)

**^13^C NMR (100 MHz, Dimethyl Sulfoxide-d6):** δ 152.91, 149.56, 144.08, 141.12, 132.54, 125.72, 125.59, 122.51, 122.09, 32.01

**HRMS (ESI) m/z:** Calculated for C_10_H_8_N_2_O_5_S+H^+^: 269.0212, found 269.0232

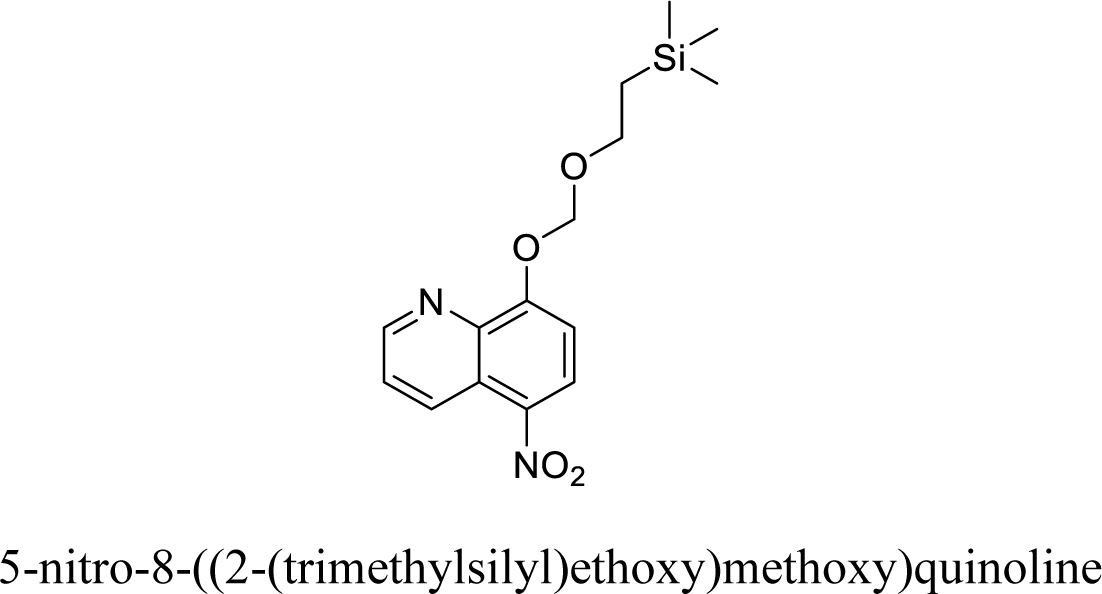

**^1^H NMR (400 MHz, Dimethyl Sulfoxide-d6):** δ 9.07 (p, J=3.84 Hz, 2H), 8.88 (d, J=4.08 Hz, 1H), 7.89 (q, J=8.72 Hz, 1H), 7.51 (d, J=8.92 Hz, 1H), 5.69 (s, 2H), 3.87 (t, J=8.24 Hz, 2H), 0.97 (t, J=8.44Hz, 2H), 0.00 (s, 9H)

**^13^C NMR (100 MHz, Dimethyl Sulfoxide-d6):** δ 159.23, 151.24, 138.89, 138.66, 132.62, 128.25, 125.70, 123.05, 110.65, 94.34, 67.52, 18.44, -0.44 ppm

**HRMS (ESI) m/z:** Calculated for C_15_H_20_N_2_O_4_Si+H^+^: 321.1265, found 321.1260

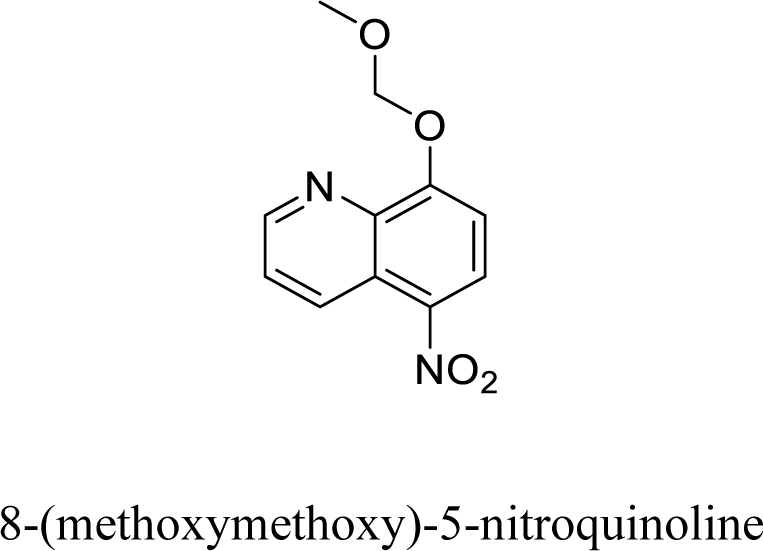

**^1^H NMR (400 MHz, Dimethyl Sulfoxide-d6):** δ 9.05-9.03 (m, 1H), 9.00 (d, J=1.66 Hz, 1H), 8.54 (d, J= 8.88 Hz 1H), 7.86 (q, J= 8.84 Hz, 1H), 7.46 (d, J= 8.88 Hz, 1H), 5.61 (s, 2H), 3.50 (s, 3H)

**^13^C NMR (100 MHz, Dimethyl Sulfoxide-d6):** δ 158.50, 150.84, 139.41, 138.39, 132.17, 127.78, 125.26, 122.59, 110.26, 95.37, 56.91

**HRMS (ESI) m/z:** Calculated for C_11_H_10_N_2_O_4_+H^+^: 235.0613, found 235.0627

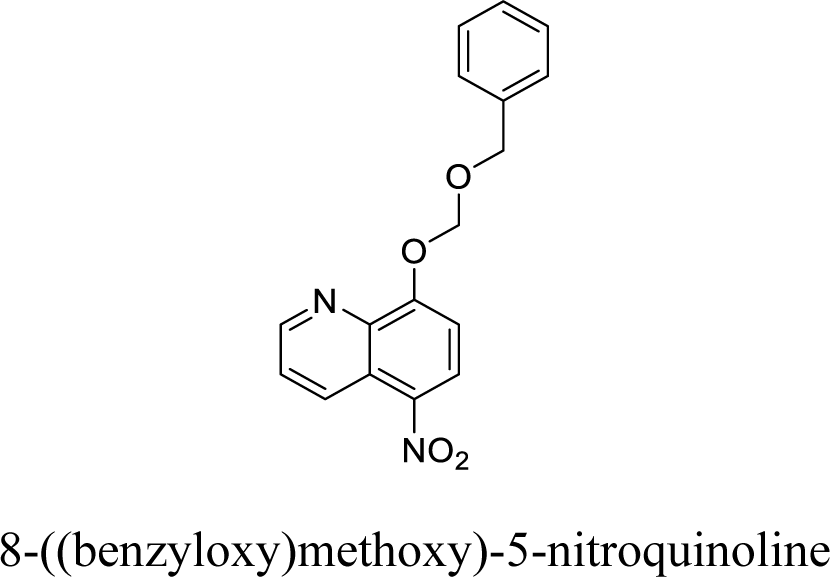

**^1^H NMR (400 MHz, Dimethyl Sulfoxide-d6):** δ 9.06-8.99 (m, 2H), 8.55 (q, J=8.60 Hz, 1H), 7.87-7.83 (m, 1H), 7.53 (q, J=8.44 Hz, 1H), 7.34-7.27 (m, 5H), 5.77 (t, J=4.44 Hz, 2H), 4.81 (d, J=7.08Hz, 2H), J= 6.92, 2H)

**^13^C NMR (100 MHz, Dimethyl Sulfoxide-d6):** δ 158.57, 150.85, 139.46, 138.41, 137.58, 132.17, 128.75, 128.27, 128.21, 127.79, 125.26, 122.60, 111.43, 93.80, 70.97ppm

**HRMS (ESI) m/z:** Calculated for C_17_H_14_N_2_O_4_+H^+^: 311.1026, found 311.1024

**<u>Compound 2</u>**

<u>^1^H-400MHz DMSO</u>

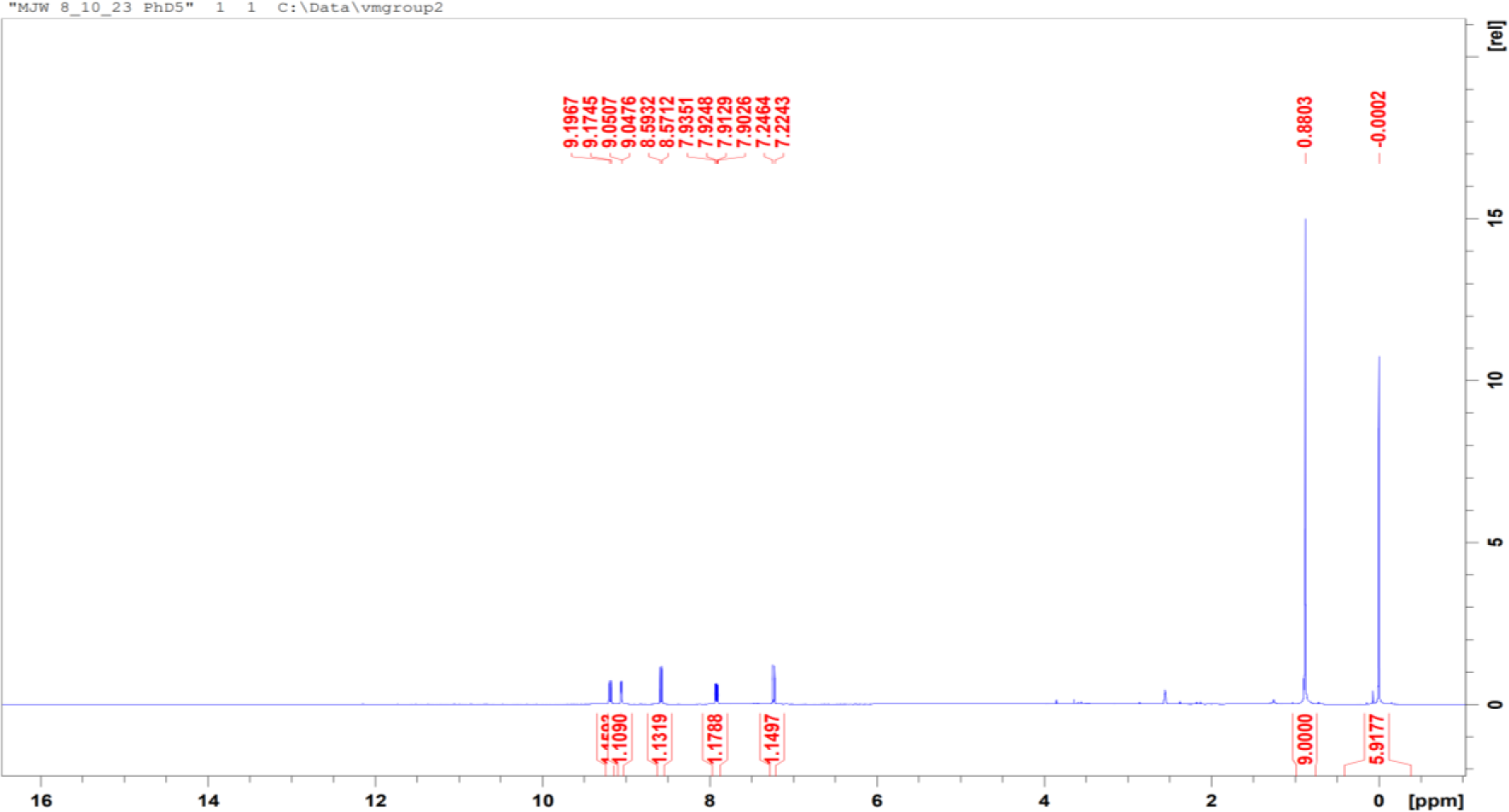

<u>^13^C-100MHz DMSO</u>

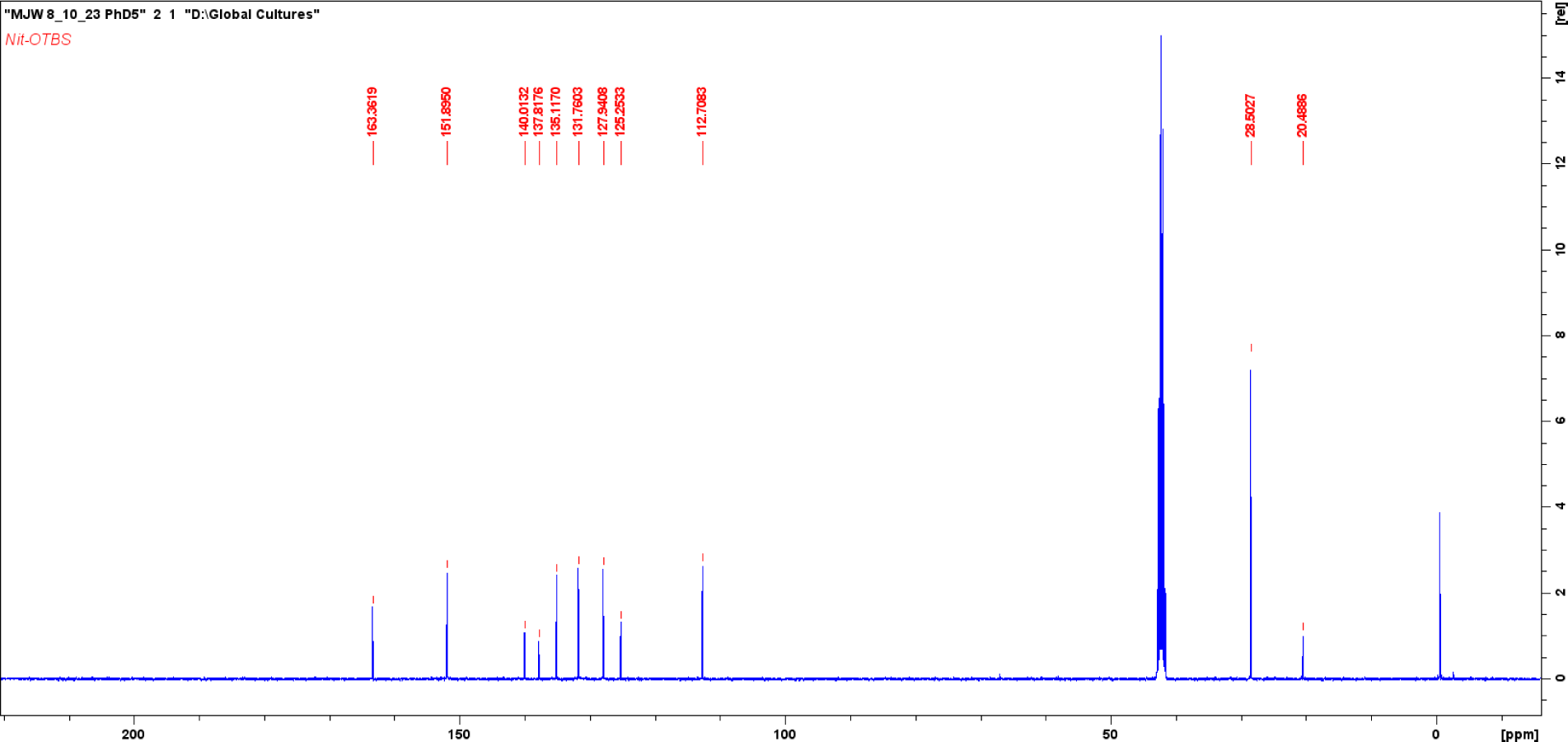

**<u>Compound 3</u>**

<u>^1^H-400MHz DMSO</u>

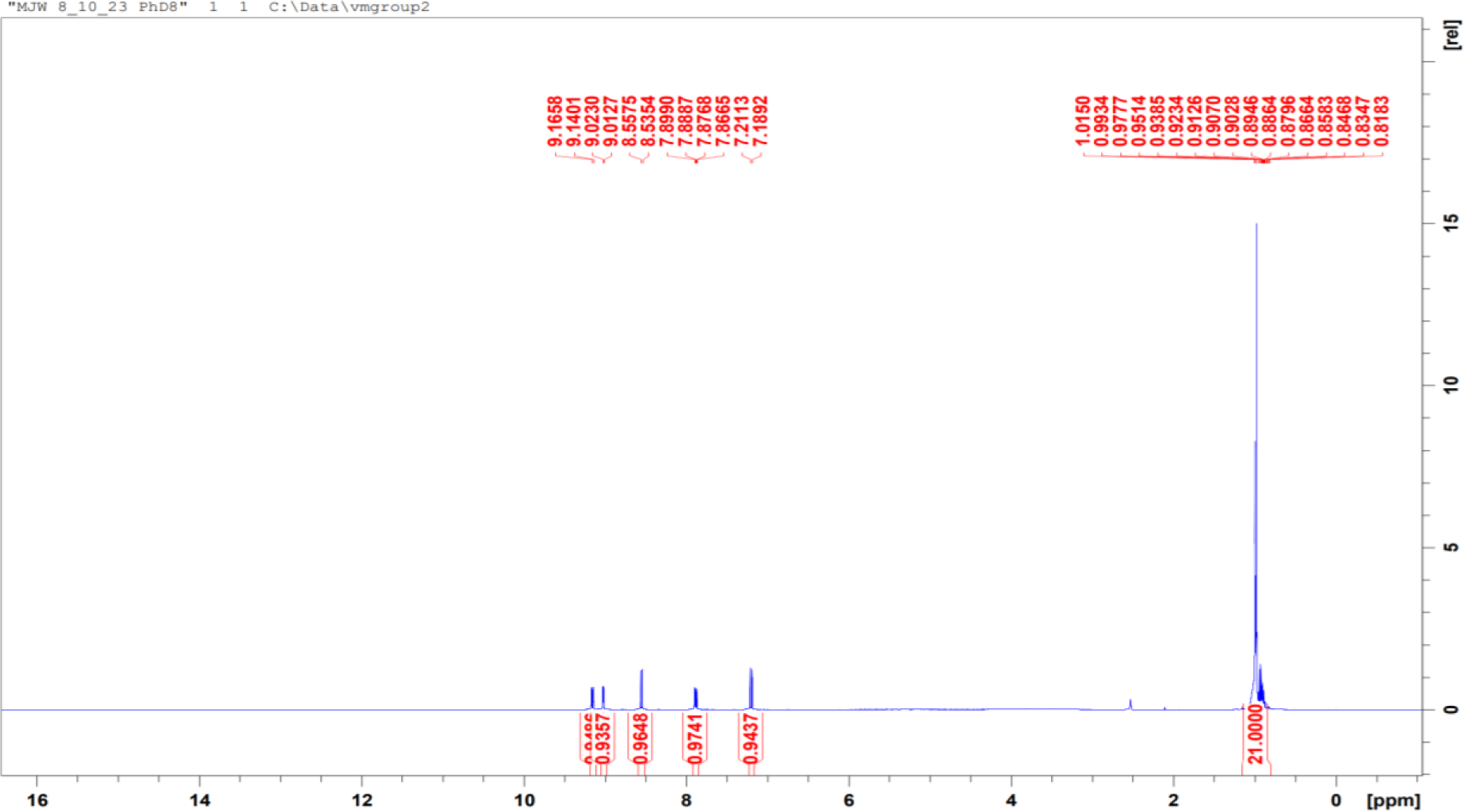

<u>^13^C-100MHz DMSO</u>

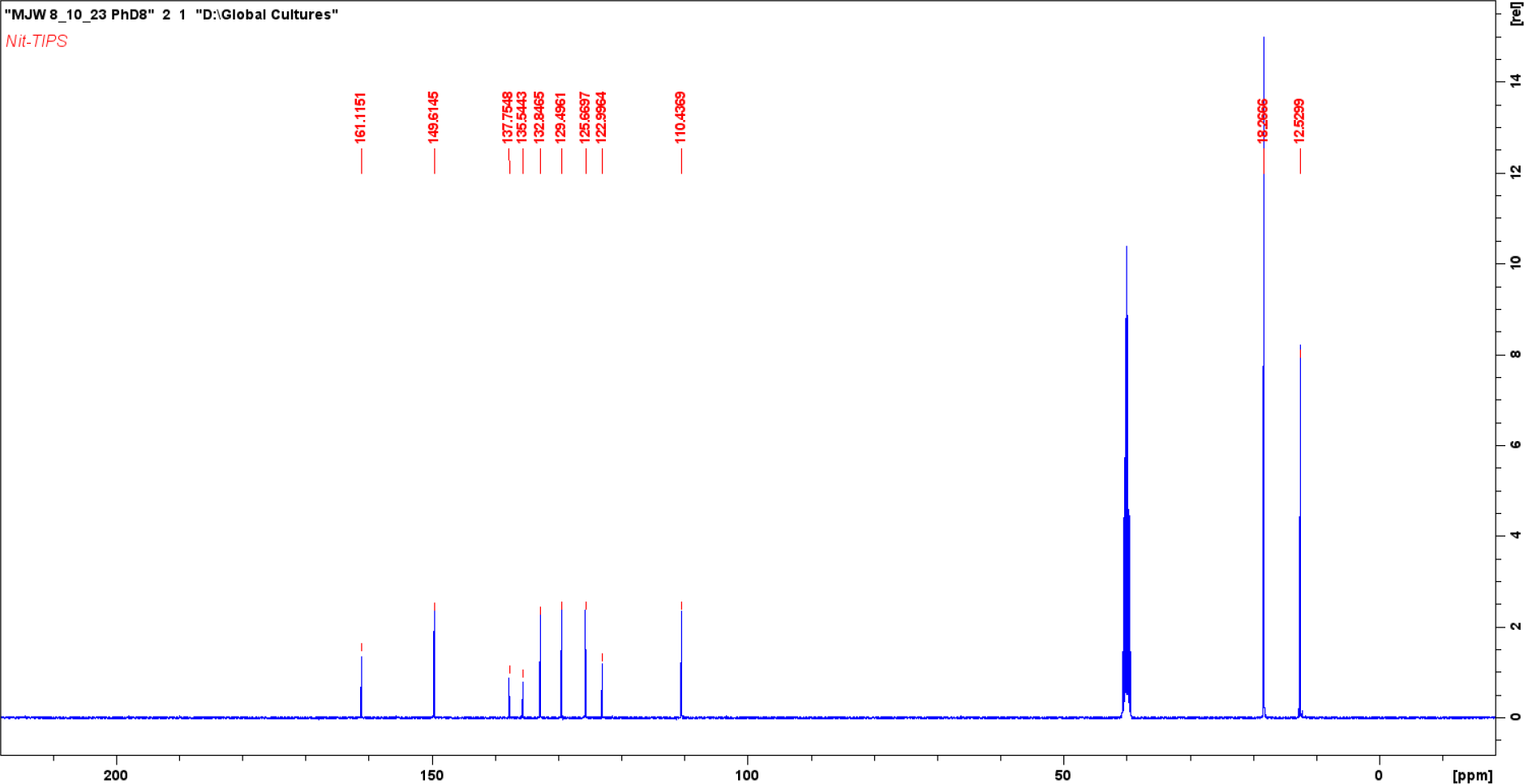

**<u>Compound 4</u>**

<u>^1^H-400MHz DMSO</u>

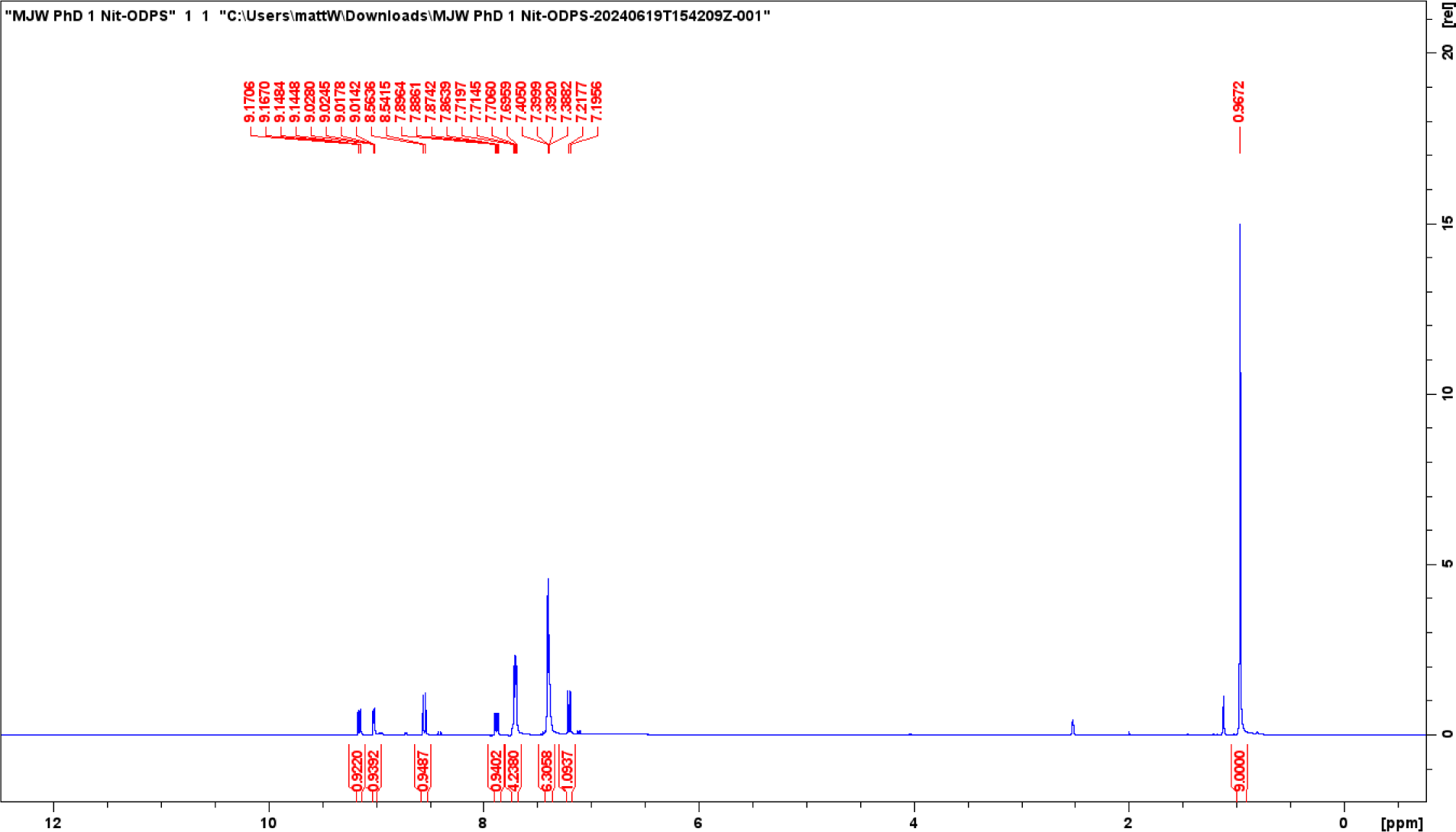

<u>^13^C-100MHz DMSO</u>

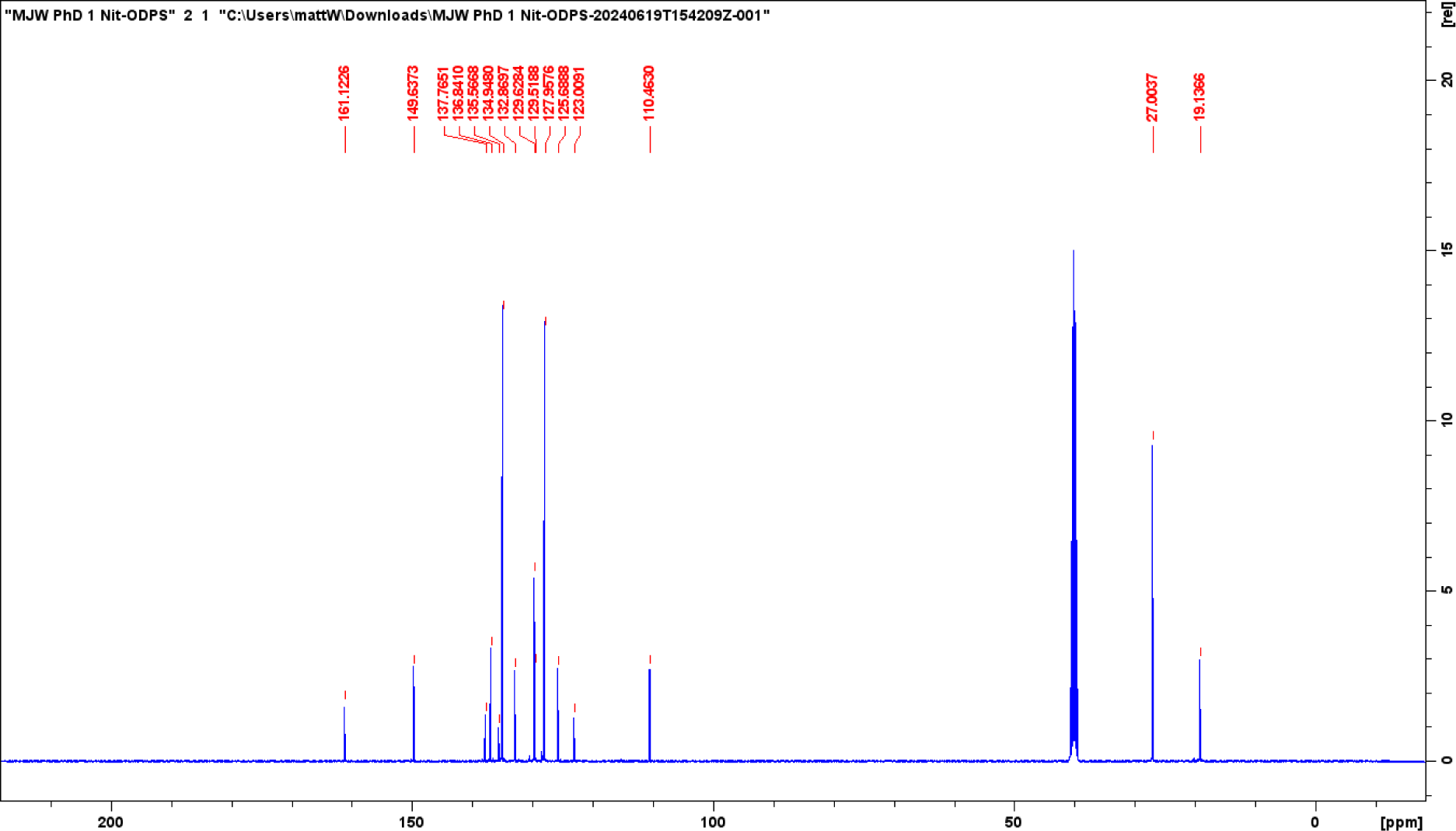

**<u>Compound 5</u>**

<u>^1^H-400MHz DMSO</u>

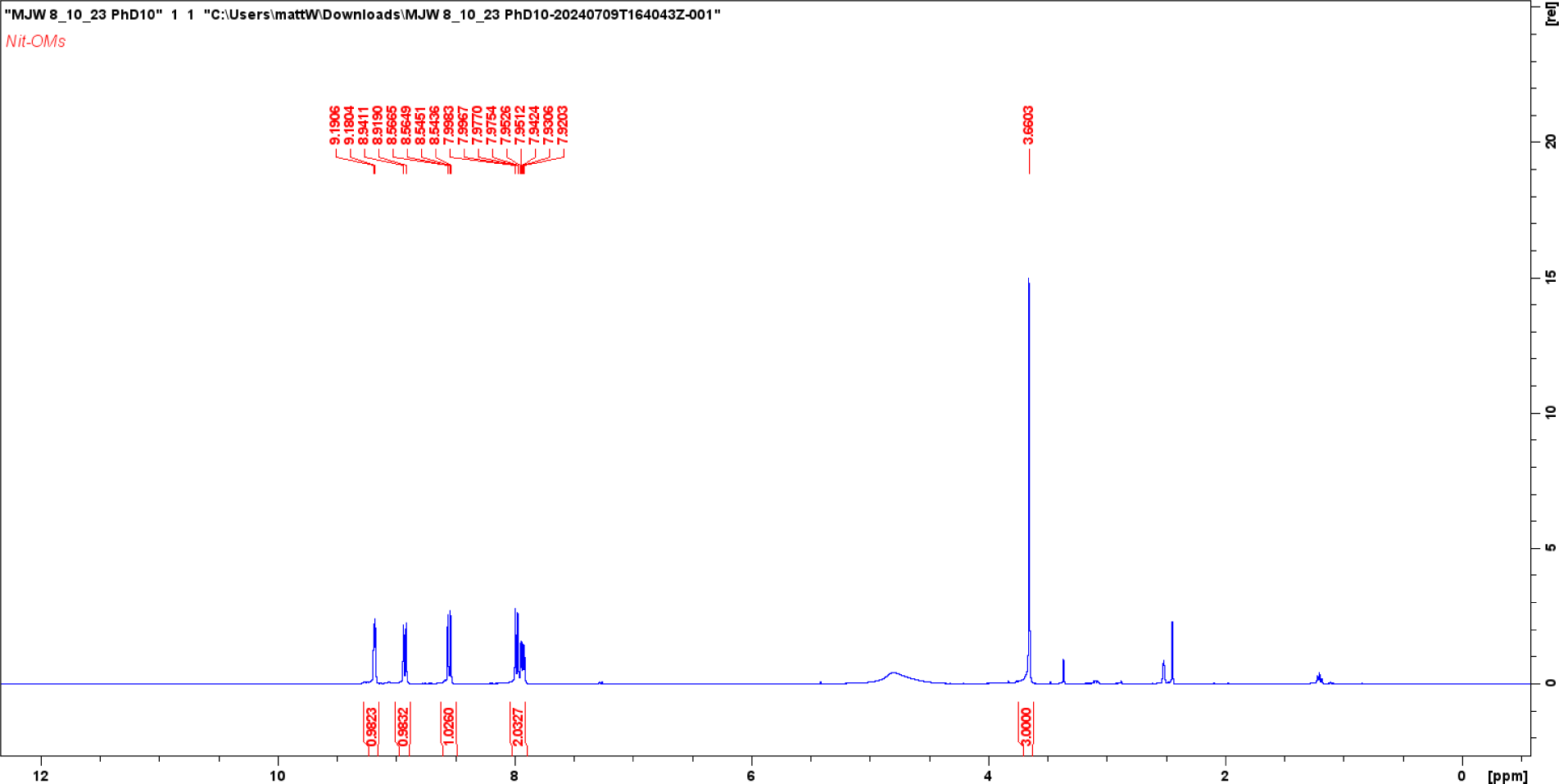

<u>^13^C-100MHz DMSO</u>

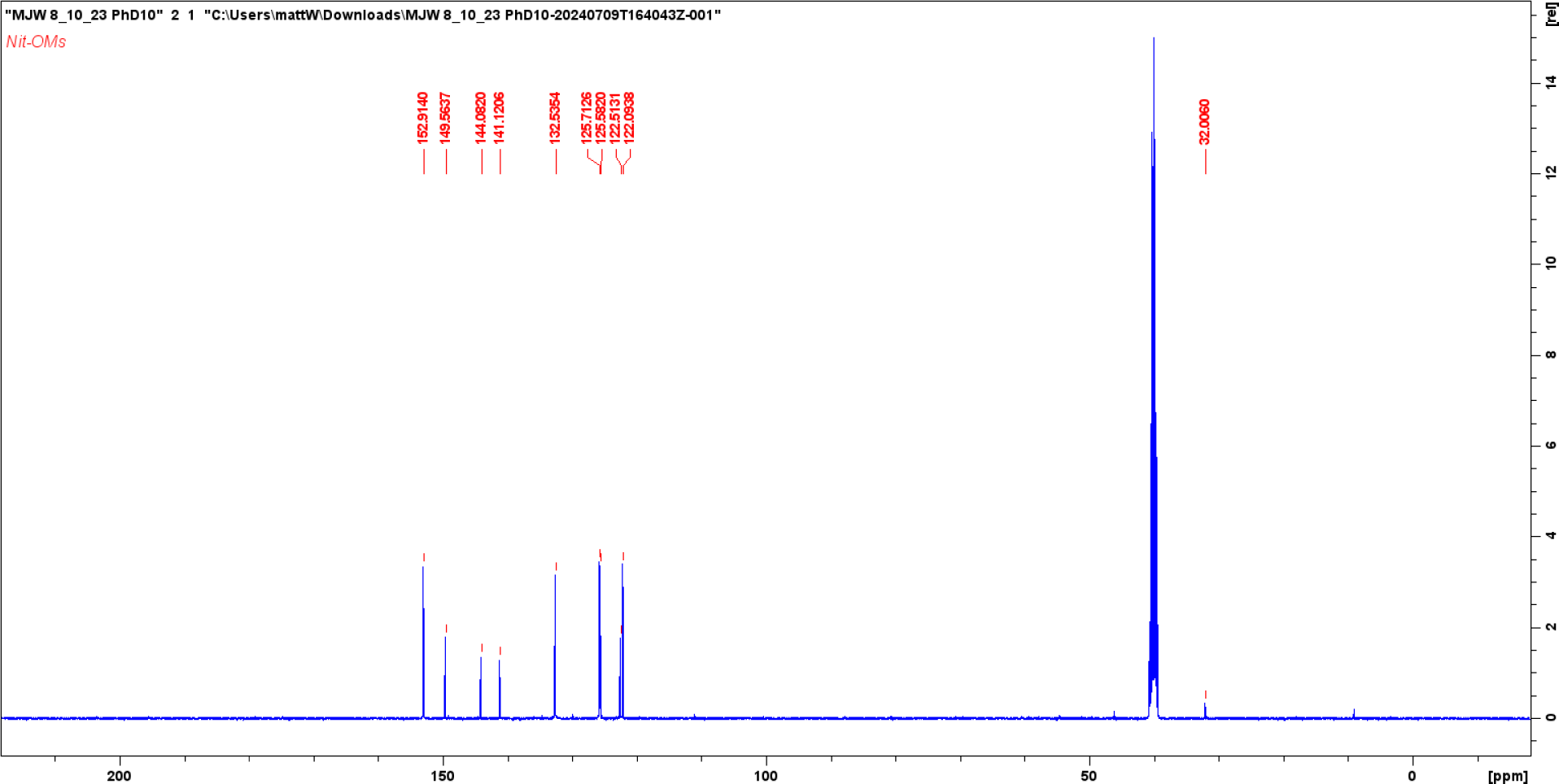

**<u>Compound 6</u>**

<u>^1^H-400MHz DMSO</u>

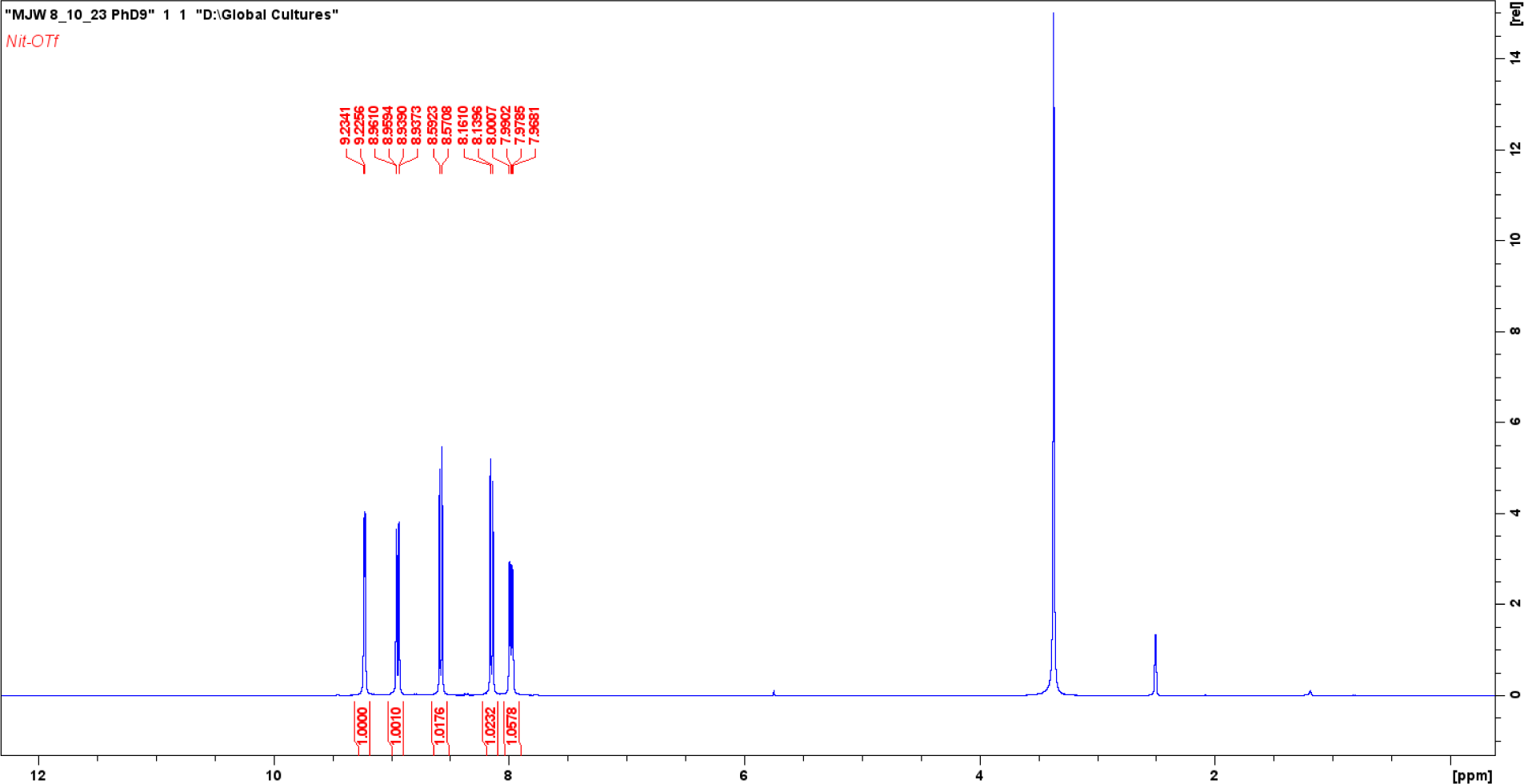

<u>^13^C-100MHz DMSO</u>

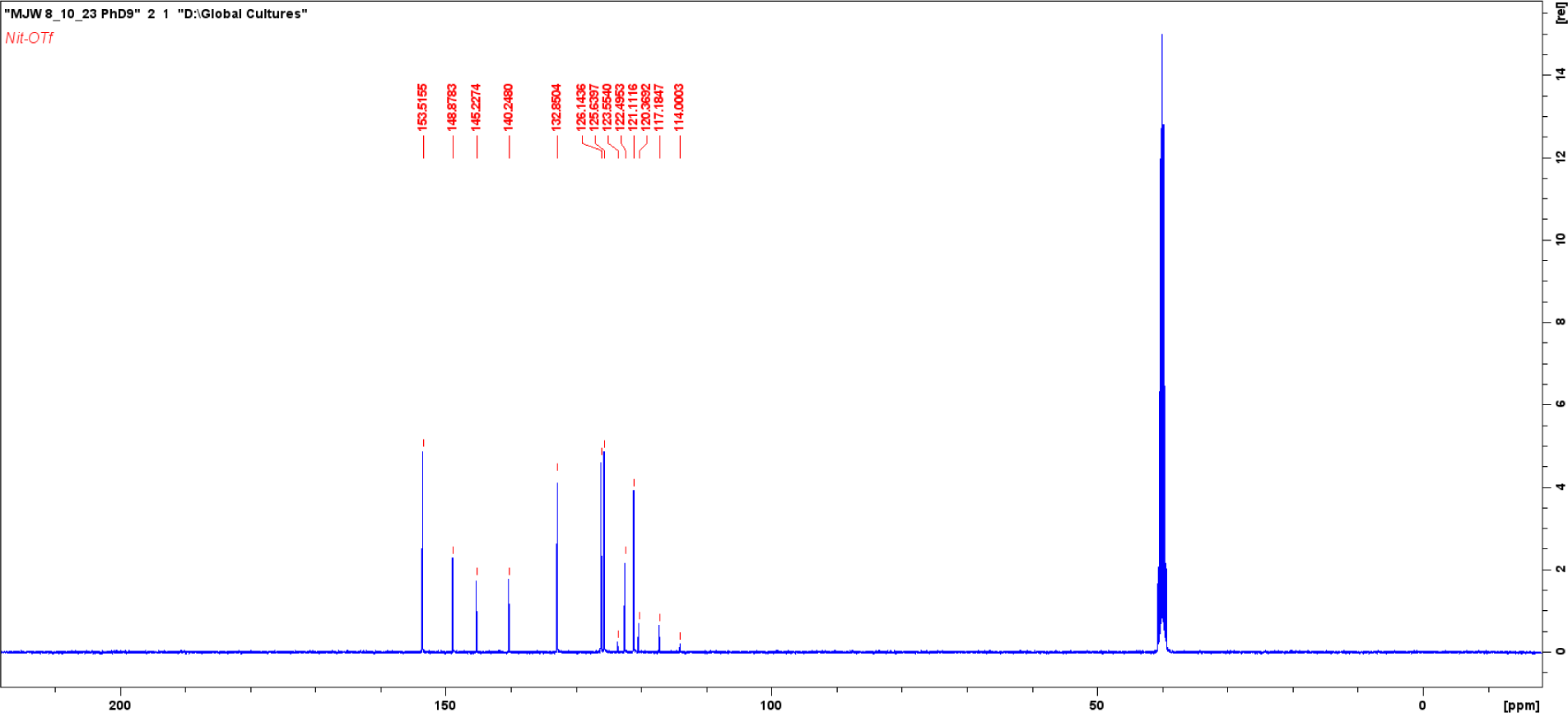

**<u>Compound 7</u>**

<u>^1^H-400MHz DMSO</u>

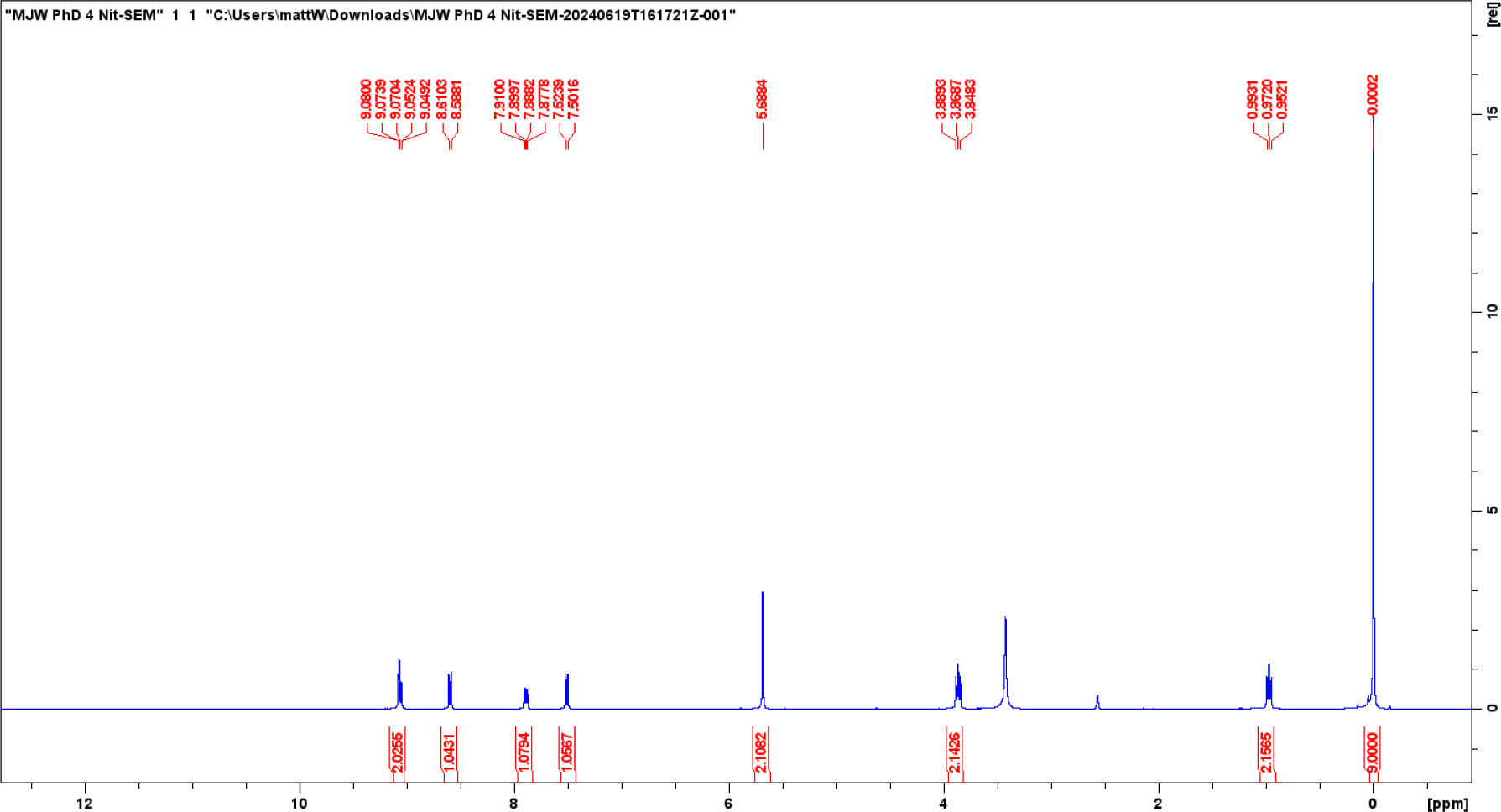

<u>^13^C-100MHz DMSO</u>

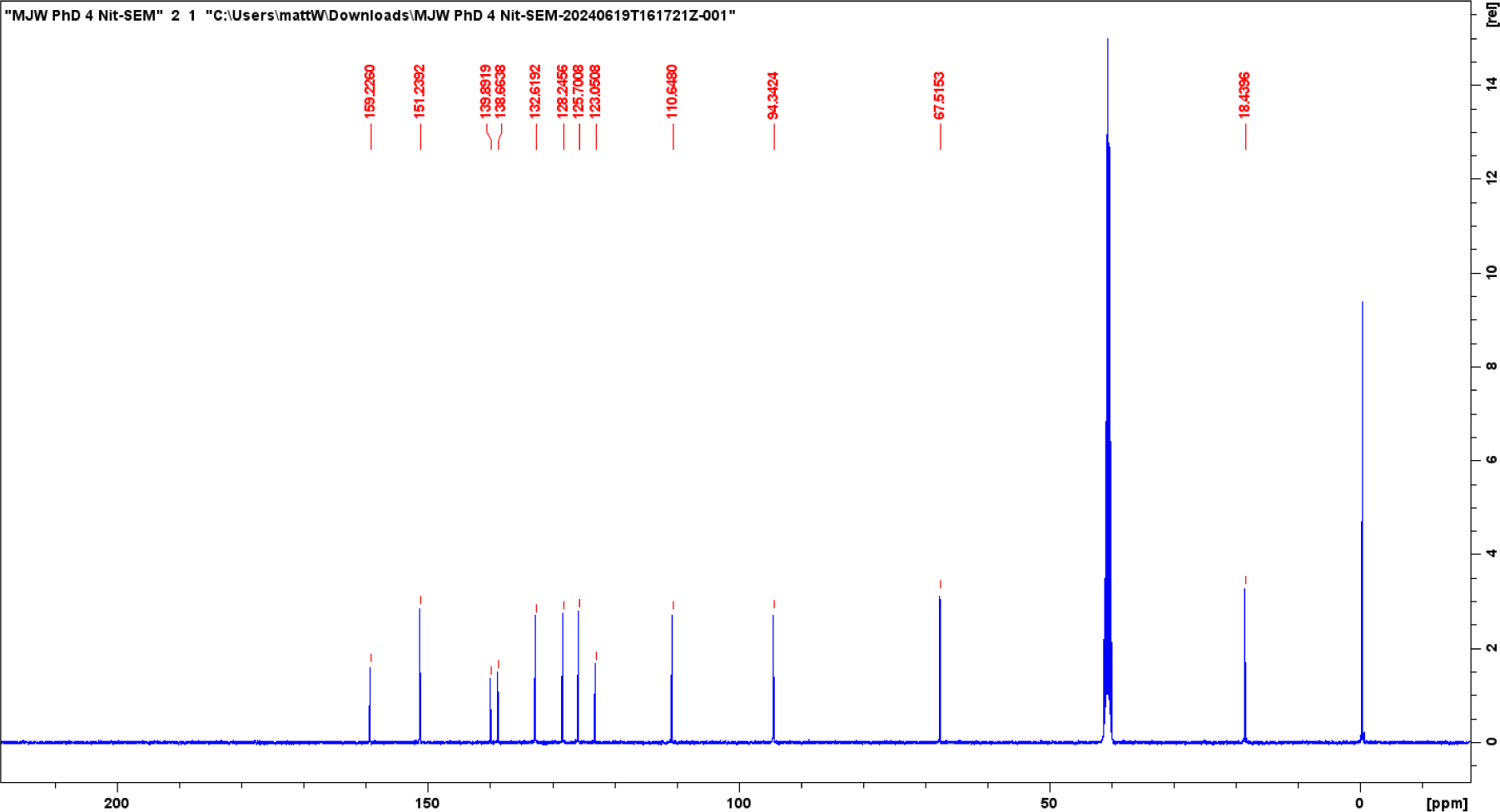

**<u>Compound 8</u>**

<u>^1^H-400MHz DMSO</u>

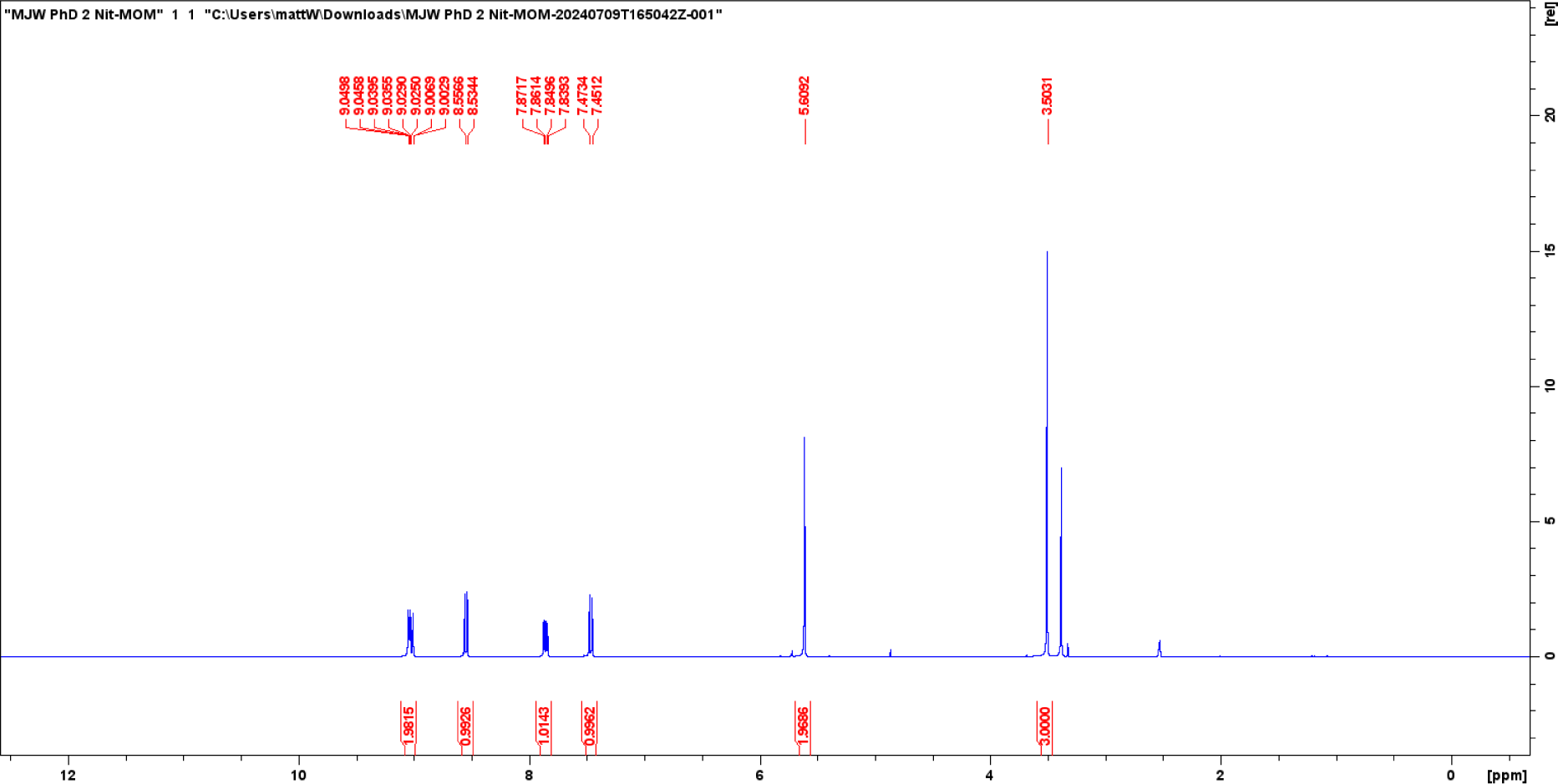

<u>^13^C-100MHz DMSO</u>

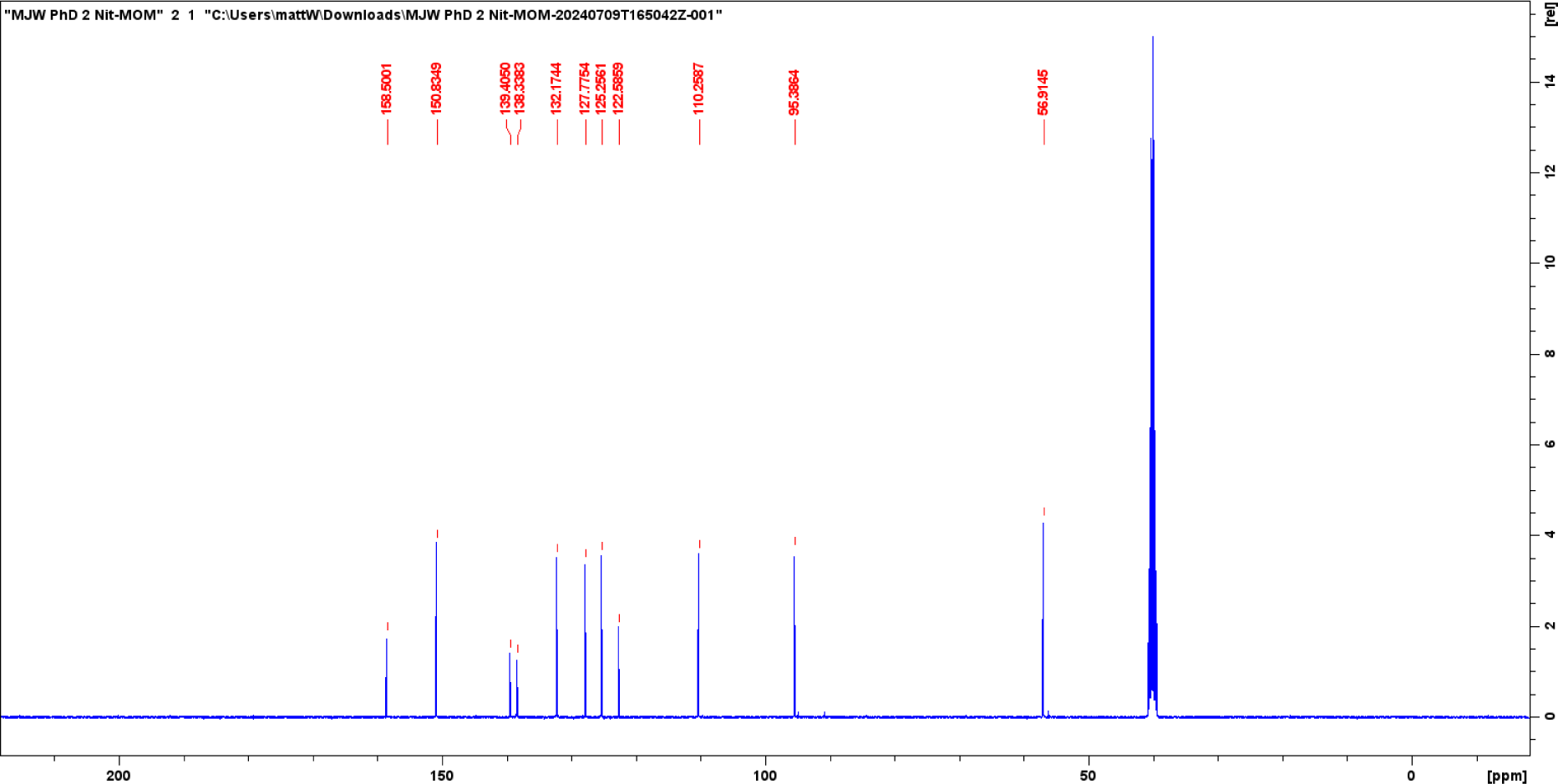

**<u>Compound 9</u>**

<u>^1^H-400MHz DMSO</u>

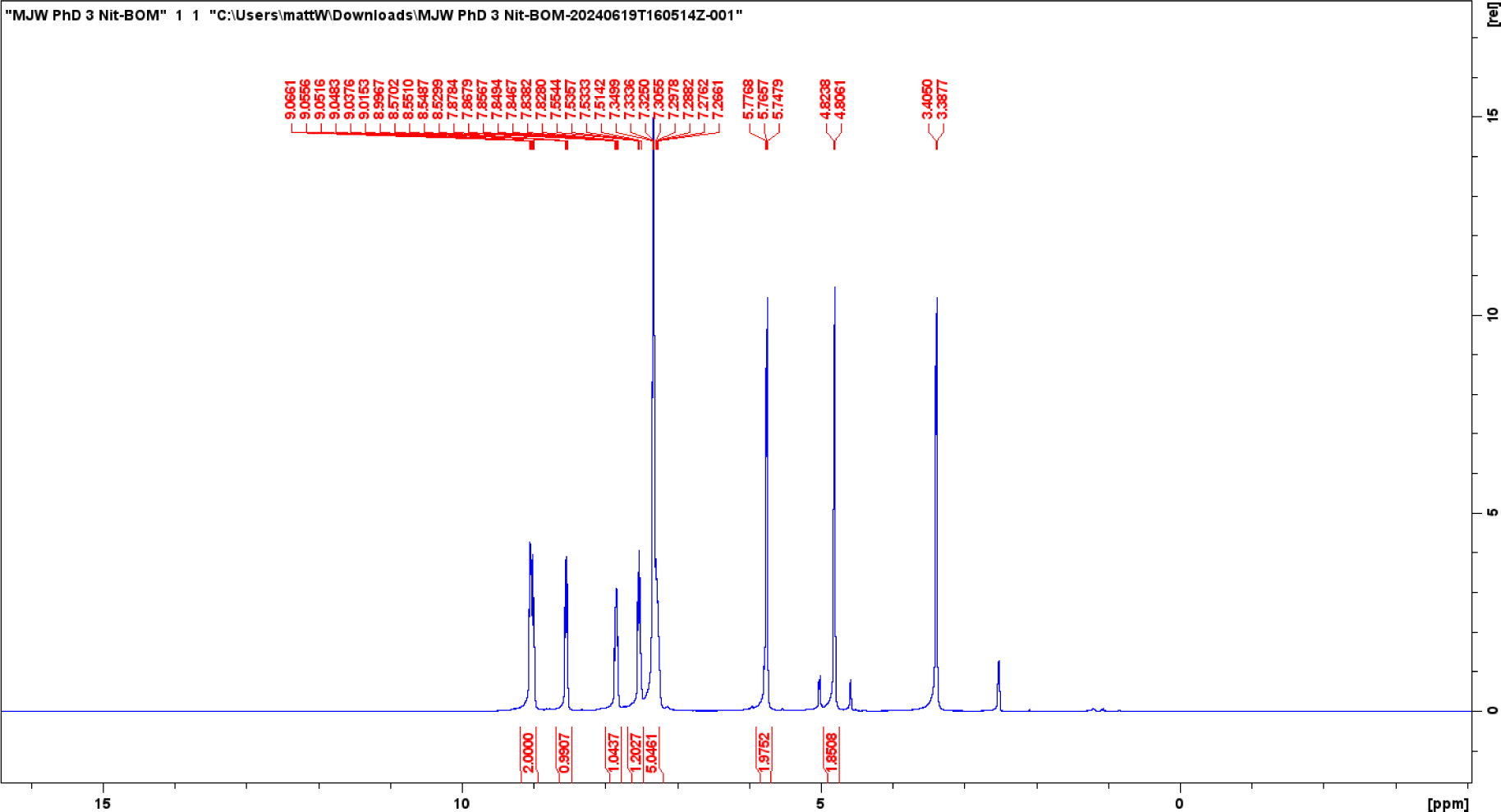

<u>^13^C-100MHz DMSO</u>

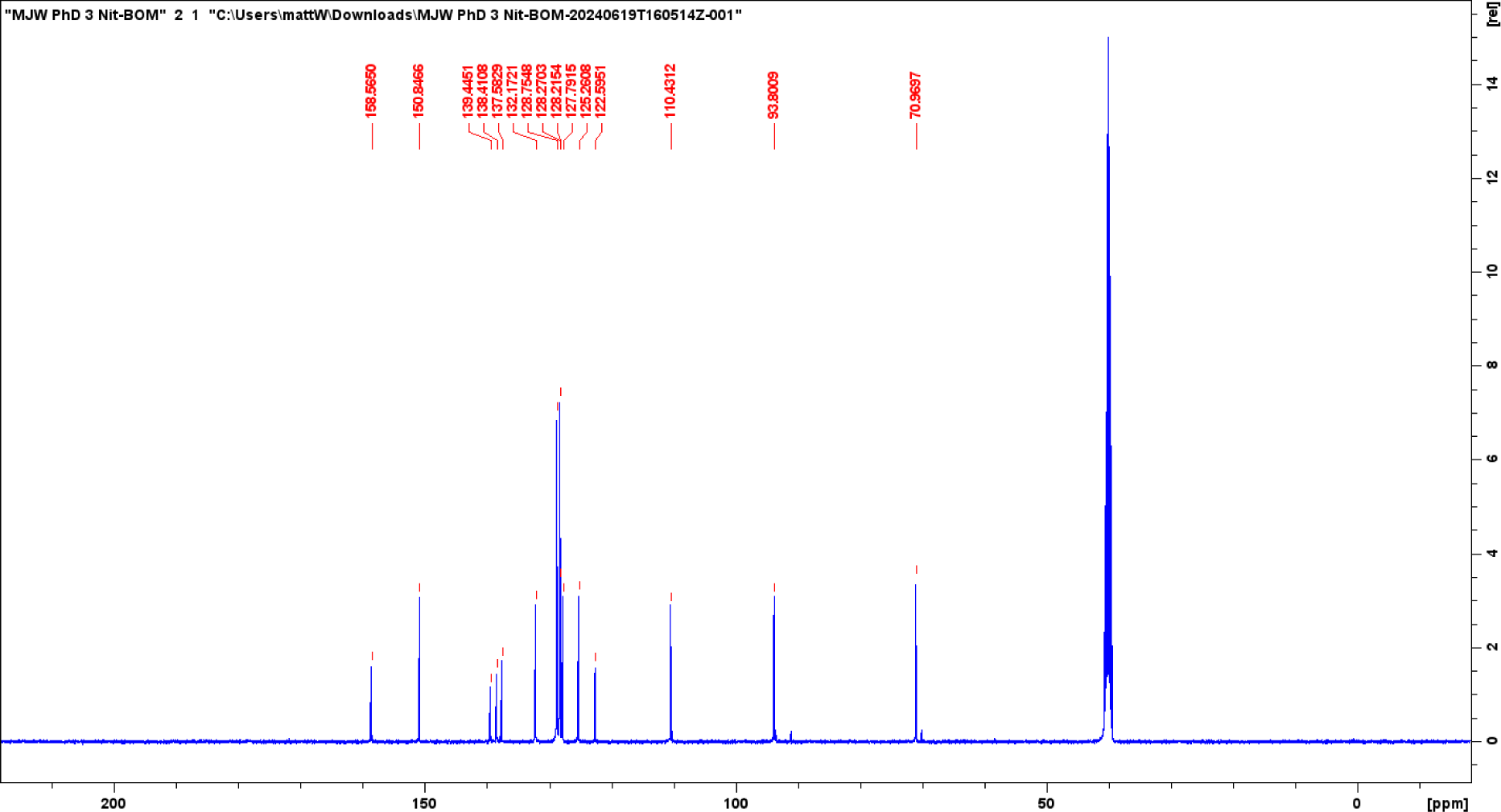

